# Novel expression of MHC II in DRG neurons attenuates paclitaxel-induced cold hypersensitivity in male and female mice

**DOI:** 10.1101/2023.03.31.535136

**Authors:** Emily E. Whitaker, Neal E. Mecum, Riley C. Cott, Diana J. Goode

## Abstract

Chemotherapy is often a life-saving treatment, but the development of intractable pain caused by chemotherapy-induced peripheral neuropathy (CIPN) is a major dose-limiting toxicity that restricts survival rates. Recent reports demonstrate that paclitaxel (PTX) robustly increases anti-inflammatory CD4^+^ T cells in the dorsal root ganglion (DRG), and that T cells and anti-inflammatory cytokines are protective against CIPN. However, the mechanism by which CD4^+^ T cells are activated, and the extent cytokines released by CD4^+^ T cells target DRG neurons are unknown. Here, we found novel expression of functional major histocompatibility complex II (MHCII) protein in DRG neurons, and CD4^+^ T cells in close proximity to DRG neurons, together suggesting CD4^+^ T cell activation and targeted cytokine release. MHCII protein is primarily expressed in small nociceptive neurons in male mouse DRG regardless of PTX, while MHCII is induced in small nociceptive neurons in female DRG after PTX. Accordingly, reducing MHCII in small nociceptive neurons increased hypersensitivity to cold only in naïve male mice, but increased severity of PTX-induced cold hypersensitivity in both sexes. Collectively, our results demonstrate expression of MHCII on DRG neurons and a functional role during homeostasis and inflammation.

**Graphical abstract:** 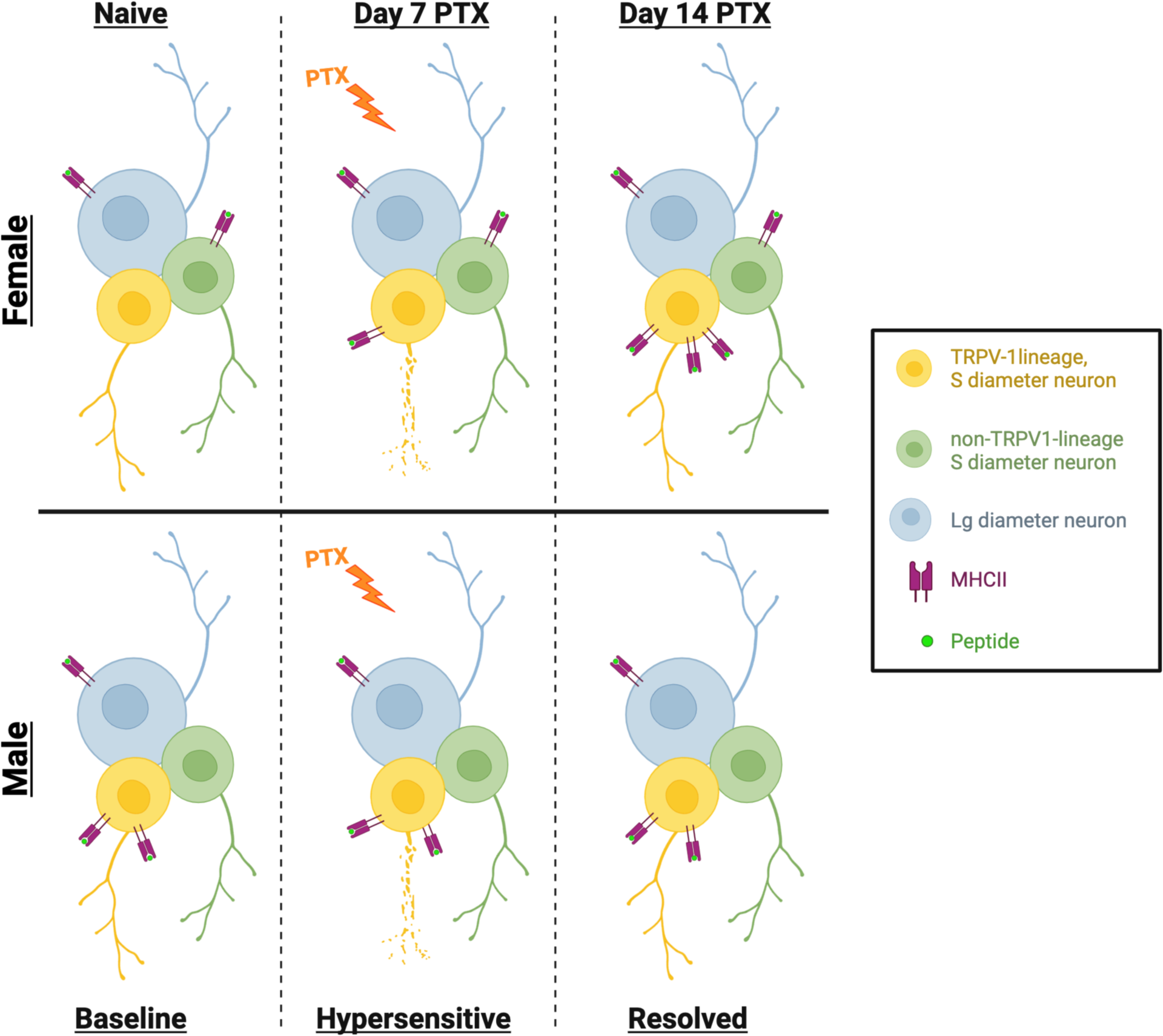

Created with Biorender.com

**Summary:** Novel expression of functional MHCII protein was detected on the surface of DRG neurons, suggesting a potential mechanism for CD4^+^ T cell activation and targeted cytokine release. Reducing MHCII from a subpopulation of neurons known to contribute to CIPN increased the severity of PTX-induced cold hypersensitivity in female and male mice.

## INTRODUCTION

Paclitaxel (PTX) is an anti-neoplastic drug commonly used to treat breast, lung, and ovarian cancer (Weaver, 2014). However, this life-saving cancer treatment is often dose-limiting as more than 60% of patients develop painful chemotherapy-induced peripheral neuropathy (CIPN) (Seretny et al., 2014), including those taking PTX (Dougherty et al., 2004). Endothelial cells encapsulating dorsal root ganglia (DRG) have large fenestrations (Jimenez-Andrade et al., 2008) with an altered repertoire of tight junction proteins (Hirakawa et al., 2004) allowing PTX to enter the DRG and to bind to toll-like receptor 4 (TLR4) expressed on macrophages and DRG neurons (Byrd-Leifer et al., 2001). Pro-inflammatory cytokines and PTX act on DRG neurons inducing hyperexcitability (Li et al., 2018, Boehmerle et al., 2006, Li et al., 2017) and neurotoxicity (Goshima et al., 2010, Flatters and Bennett, 2006), which manifests as pain, tingling, and numbness in a stocking and glove distribution (Rowinsky et al., 1993). Recently, T cells and anti-inflammatory cytokines (IL-4 and IL-10) have been shown to suppress CIPN (Krukowski et al., 2016, Laumet et al., 2020, Chen et al., 2020); however, the underlying mechanism contributing to the resolution of CIPN remains elusive, which is evidenced by limited treatment options.

While treatment with PTX robustly increases anti-inflammatory CD4^+^ T cells in female mouse DRG (Goode et al., 2022), the mechanism by which this occurs is unclear. It is thought that communication between CD4^+^ T cells and neurons primarily occurs through the release of soluble mediators as neurons express cytokine receptors (Copray et al., 2001) while CD4^+^ T cells express receptors for neurotransmitters (Wang et al., 1992). Pro-inflammatory CD4^+^ T cell subsets (Th1 and Th17) secrete cytokines that sensitize neurons in the DRG and spinal cord to increase nociceptive-signaling (Richter et al., 2012, Vikman et al., 2003) while anti-inflammatory subsets (FoxP3 Tregs and Th2) secrete IL-10 and IL-4, which suppress activity in sensitized nociceptors and reduce neuronal hyperexcitability (Krukowski et al., 2016), respectively. Although CD4^+^ T cell–neuron communication through soluble mediators has been more extensively studied, published DRG neuron RNA seq mouse and human datasets (Nguyen et al., 2021, Tavares-Ferreira et al., 2022) indicate direct cell-cell interaction is possible. By mining these single cell RNA seq data sets, we found that DRG neurons express major histocompatibility complex II (MHCII) transcripts (Usoskin et al., 2015, Ray et al., 2018), but there are no reports that show MHCII protein or function in DRG neurons.

MHCII is traditionally thought to be constitutively expressed on antigen-presenting cells (APCs) and induced by inflammation on some non-APCs, including endothelial, epithelial, and glial cells (van Velzen et al., 2009). In contrast to these non-APCs, human neural stem cells (hNSCs) and neural progenitor cells within embryo spinal cord and DRG constitutively express MHCII protein prior to the development of the adaptive immune response (Vagaska et al., 2016). Differentiating hNSCs to neurons in vitro increased MHCII with inflammatory stimuli (IFN-γ) further elevating expression (Vagaska et al., 2016). Finding MHCII expression on terminally differentiated neurons in vivo would provide insight into the mechanism by which CD4^+^ T cells contribute to pain and neurological diseases.

While DRG neurons are wrapped with satellite glial cells (SGCs), natural gaps in the glial envelope can occur (Dixon, 1969). Furthermore, sciatic nerve injury and nerve transection models demonstrate that immune cells can breach the SGC and lie directly against the neuron (Hu and McLachlan, 2002), suggesting direct neuron-immune communication. In our present study, we found that CD4^+^ T cells can breach the SGC barrier during homeostasis and inflammatory conditions and that DRG neurons express functional surface-bound MHCII protein. Moreover, neuronal MHCII represents a novel mechanism to suppress CIPN, which could be exploited for therapeutic intervention against not only pain but potentially autoimmunity and neurological diseases.

## RESULTS

### CD4^+^ T cells breach the SGC barrier in mouse DRG

We recently demonstrated that CD4^+^ T cells in the DRG primarily produce anti-inflammatory cytokines (Goode et al., 2022). Due to their short half-life, cytokine release typically occurs in close proximity to the target cell; however, the extent cytokines released by CD4^+^ T cells target DRG neurons in vivo is unknown. Therefore, initial fluorescent imaging experiments were designed to visualize CD4^+^ T cell proximity to neurons in DRG tissue. To increase the likelihood of detecting CD4^+^ T cells, we utilized female mice as we have previously shown that the frequency of CD4^+^ T cells in female mouse DRG is higher than male (Goode et al., 2022). We found that CD4^+^ T cells were consistently present in female DRG (19.6 CD4^+^ T cells/mm^2^ for naïve **(Fig. S1)** and 38.4 for day 14 PTX-treated female mice (**Fig. 1A**), n=7/condition) and frequently clustered in DRG tissue in close proximity to neurons. Confocal microscopy was required to demonstrate that CD4^+^ T cells colocalized with neurons in the absence of SGC markers (GLAST/FABP7) in DRG tissue from naïve and day 14 PTX-treated female mice (**Fig. 1B-D, S2 3D video, S3)**. Z-stacks of DRG tissue confirmed that CD4^+^ T cells were located between the border (identified by differential interference contrast microscopy) of DRG neurons and SGCs (**Fig. 1B, C, S3**). Our results demonstrate that CD4^+^ T cells can breach the SGCs barrier, suggesting CD4^+^ T cell– DRG neuron communication can occur through both cell-cell contact and the release of soluble mediators.

**Fig. 1.**
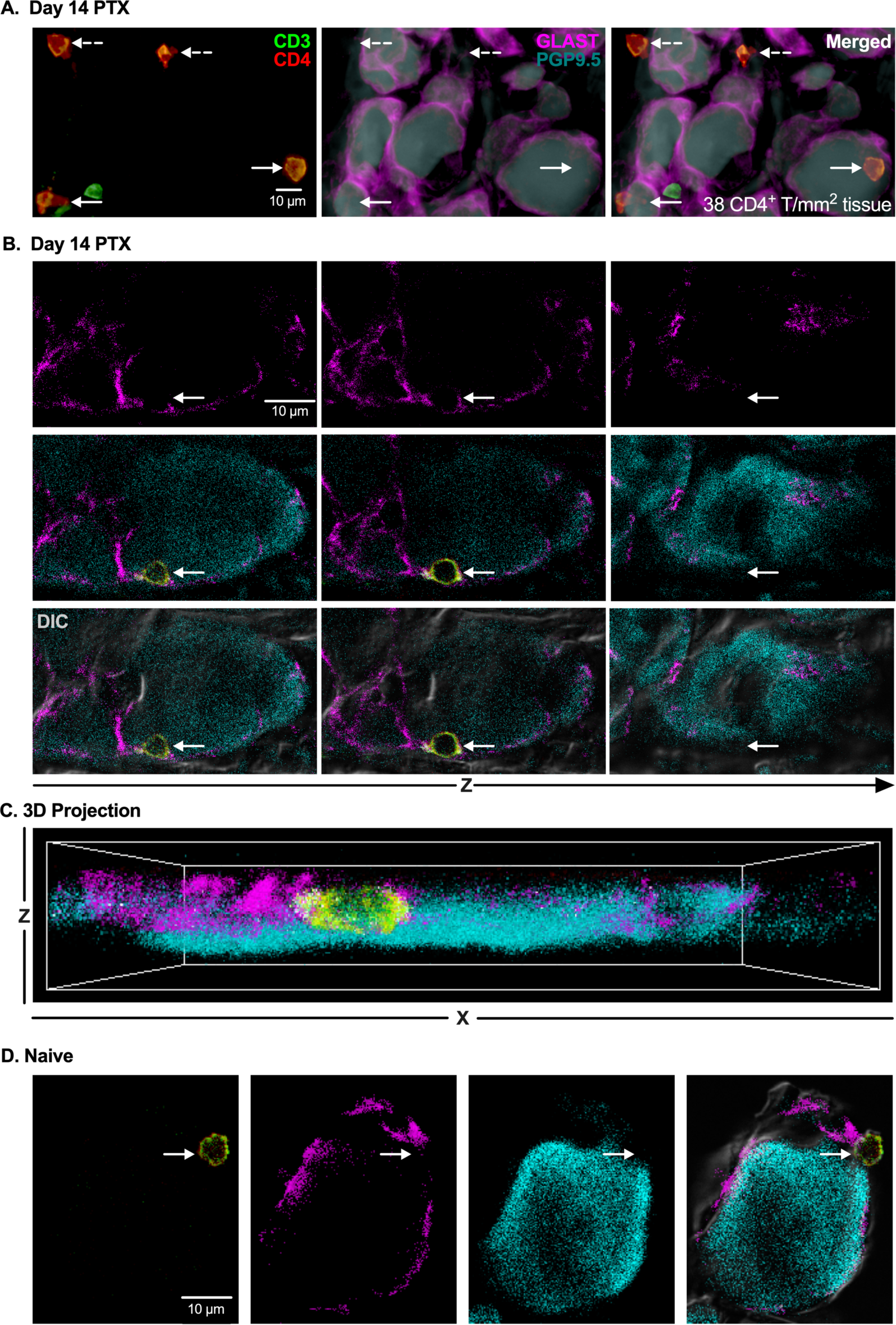
CD4^+^ T cells breach SGC barrier surrounding neurons in DRG tissue from naïve and PTX-treated female mice. **(A-D)** Immunohistochemical (IHC) staining of neurons (PGP9.5: cyan), SGCs (GLAST: magenta), and T cells (CD3: green, CD4: red) in L4 DRG tissue from naïve and day 14 PTX-treated female mice. **(A)** Representative widefield fluorescence two-channel (CD3/CD4; GLAST/PGP9.5) and merged images of CD4^+^ T cells and DRG neurons in close proximity (solid arrow = absence of GLAST; dashed arrow = occluded by GLAST; n=5) in PTX-treated female tissue. **(B)** Differential inference contrast (DIC), fluorescence, and merged confocal Z-slice images of a CD4^+^ T cell between the SGC barrier and a PGP9.5^+^ DRG neuron from a PTX-treated mouse. **(C)** 3D projection of CD4^+^ T cell shown in **(B and video in Fig. S2)**. **(D)** Maximum projection images of a CD4^+^ T cell breaching the SGC barrier surrounding a DRG neuron from a naïve mouse.

Expression of immune molecules on DRG neurons could facilitate communication between CD4^+^ T cells and DRG neurons. Indeed, published RNA seq data sets demonstrate that both human and mouse DRG neurons (Usoskin et al., 2015, Lopes et al., 2017) express MHCII and MHCII-associated genes, which are critical for the activation of CD4^+^ T cells upon cell-cell contact. Consistent with the RNA seq data, which shows expression of RFX1 RNA (!!! INVALID CITATION !!! (Usoskin et al., 2015, Lopes et al., 2017)), we detected RFX1 protein, a transcription factor for MHCII, in the nucleus of DRG neurons in male and female mice **(Fig. S4)**, further supporting the possibility of MHCII protein in DRG neurons.

### MHCII protein detected in DRG neurons

Inflammatory stimuli are known to increase the expression of MHCII in immune cells (Pfannenstiel et al., 2010); therefore, female mice were injected with PTX to induce inflammation in the DRG. DRG neuron lysates from naïve and day 14 PTX-treated female and male mice were probed with an antibody against MHCII and analyzed by western blot. Consistent with inflammatory stimuli boosting the expression of MHCII, DRG neurons from PTX-treated female mice had almost 2-fold more MHCII protein than DRG neurons from naïve female mice (29.78 ± 2.76 arbitrary fluorescent units (AFUs) from naïve to 52.23 AFUs ± 8.21 14 days post-PTX, normalized to tubulin, p=0.0410, n=4/condition) (**Fig. 2A, B**). In contrast, PTX treatment did not change the amount of MHCII protein detected in DRG neurons from male mice (30.97 ± 2.68 AFUs from naïve to 27.12 AFUs ± 8.91 14 days post-PTX, p=0.9377, n=4/condition).

**Fig. 2.**
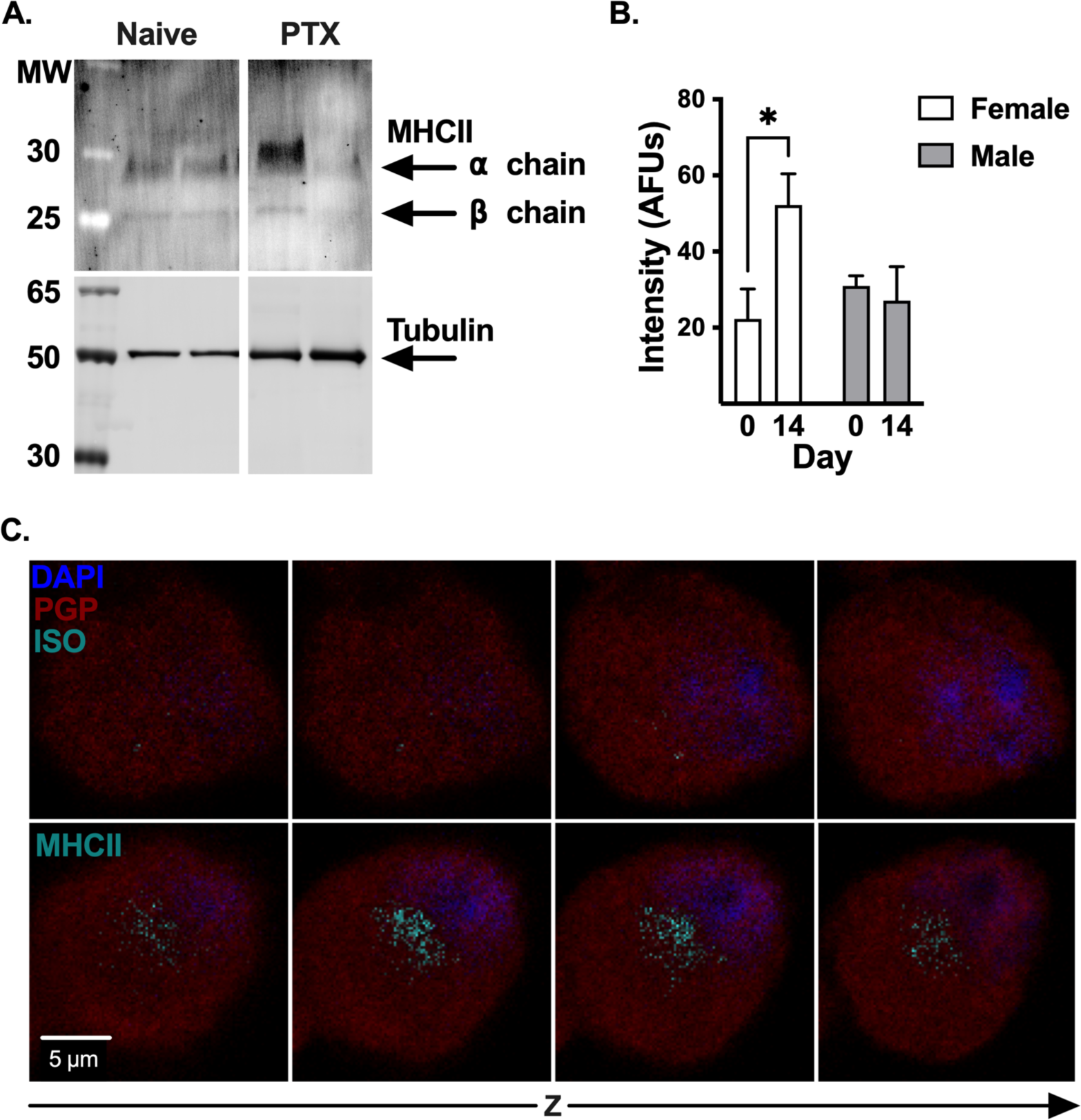
DRG neurons from female and male mice express MHCII protein. **(A)** Western blot of MACS-DRG neuron lysates from naïve and day 14 PTX-treated female and male mice, probed with antibodies against MHCII and beta tubulin (loading control). **(B)** Quantification of MHCII band intensity normalized to tubulin for DRG neuron lysates from naïve and day 14 PTX-treated female (white) and male (grey) mice shown in **(A)**. Statistical significance determined by 2-way ANOVA with Sidak’s multiple comparison test (*p <0.05, n=4/condition). **(C)** Confocal stepwise Z-slices of cultured PGP9.5^+^ DRG neurons (red) from day 14 PTX-treated female mice stained with DAPI (blue) and MHCII or isotype-control antibody (cyan).

Although cell lysates used for western blot primarily consisted of DRG neurons, negative selection by magnetic-activated cell sorting (MACS) does not yield a cell sample that is 100% pure. Therefore, it is possible that the MHCII protein signal detected by western blot represents MHCII from contaminating non-neuronal cells. To verify that DRG neurons express MHCII protein, we visualized MHCII by immunocytochemistry (ICC) of cultured DRG neurons from day 14 PTX-treated female mice, which had the highest expression of MHCII by western blot. Nuclei were labeled with DAPI to eliminate the possibility that MHCII staining was the result of a contaminating non-neuronal MHCII^+^ cell. Z-stack images acquired by confocal microscopy demonstrate MHCII staining through the entire DRG neuron from day 14 PTX-treated female mice compared to the isotype-stained control (**Fig. 2C**), confirming expression of MHCII protein in DRG neurons.

### DRG neurons express surface-MHCII

While endosomal MHCII can promote TLR signaling events(Liu et al., 2011), MHCII on the cell surface is required to activate CD4^+^ T cells. We performed flow cytometry to determine whether neurons express surface-MHCII. PGP9.5^+^ neurons comprised 28.93% ± 2.33 of total DRG cells **(Fig. S5)**, and within the PGP9.5^+^ neuron gate, 9.66% ± 0.64 of cells from naïve female mice expressed surface-MHCII (**Fig. 3A, B**). Surface-MHCII on DRG neurons increased to 15.44% ± 0.84 after treatment with PTX (p<0.0001, n=9) (**Fig. 3A, B**), which is consistent with inflammation increasing the half-life of surface-MHCII on APCs (Cella et al., 1997). In agreement with the western blot, surface-MHCII did not increase after PTX on DRG neurons from male mice (13.85% ± 1.23 in naïve and 12.10% ± 1.04 14 days post-PTX, p=0.5271, n=3-6) (**Fig. 3B**). MHCII on the neuron surface suggests a mechanism by which CD4^+^ T cells could directly communicate with DRG neurons.

**Fig. 3.**
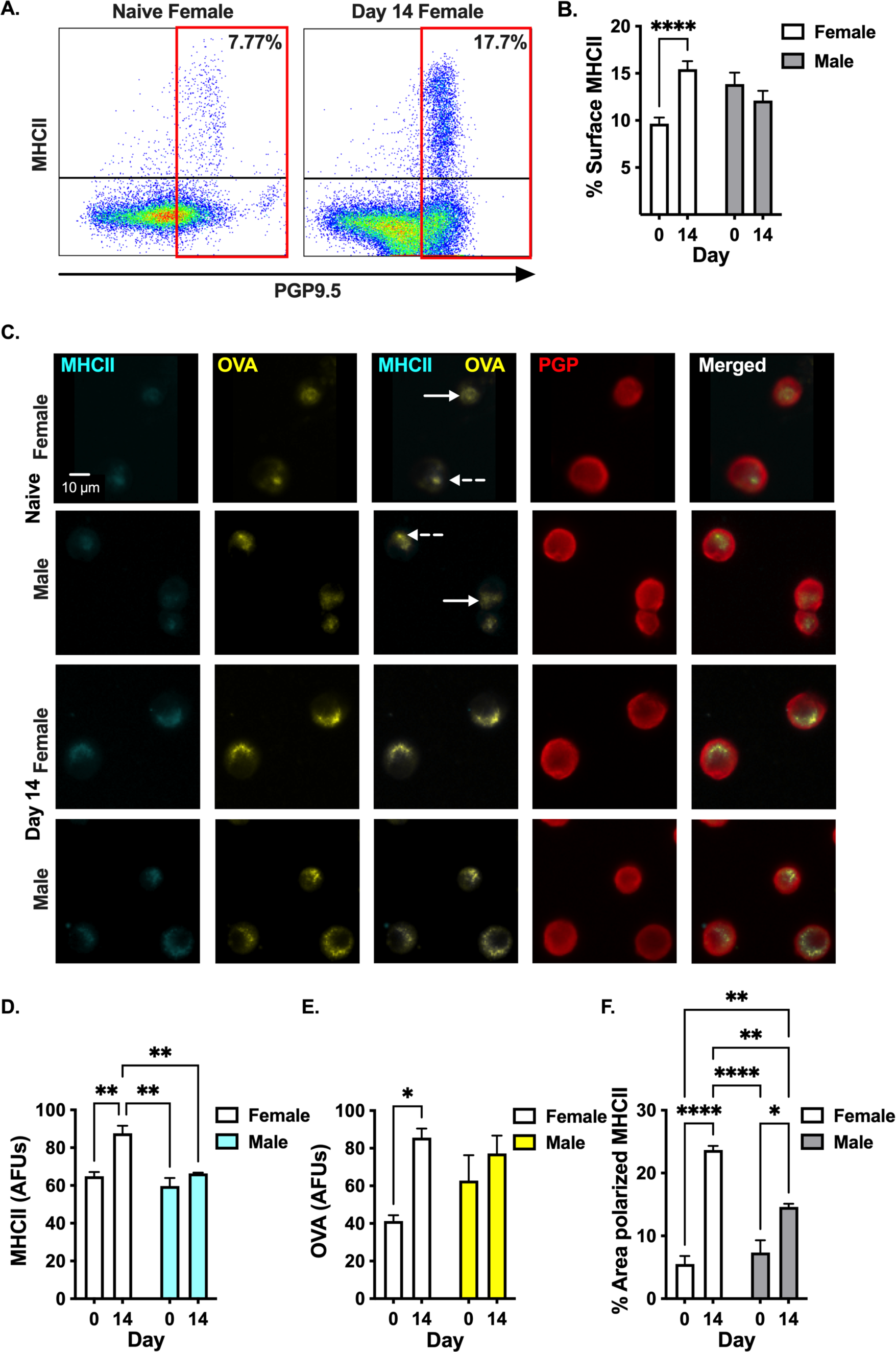
PTX treatment increases surface expression of peptide-bound MHCII on DRG neurons from female mice. **(A)** Representative flow cytometric dot plots of the frequency of DRG neurons expressing surface-MHCII from naïve and day 14 PTX-treated female mice. **(B)** Quantification of the frequency of DRG neurons expressing surface-MHCII shown in **(A)**. **(C)** Representative widefield fluorescence microscopy images of DRG neurons (PGP9.5^+^: red) expressing surface-MHCII (cyan) co-localized with OVA-FITC (yellow) from naïve and day 14 PTX-treated female and male mice. Diffuse (solid arrow) and punctate (dashed arrow) surface-MHCII staining in DRG neurons from naïve female and male mice. Quantification of **(D)** MHCII intensity, **(E)** OVA intensity, and **(F)** percent of neuron area with polarized MHCII from **(C)**. Statistical significance determined by 2-way ANOVA with Sidak’s multiple comparison test (*p <0.05, **p < 0.01, ***p < 0.001, n=3/treatment).

CD4^+^ T cells respond to specific peptide bound to surface-MHCII; however, it is unknown if surface-MHCII on neurons can present peptide. To address this question, we cultured DRG neurons from naïve and day 14 PTX-treated female and male mice with FITC-conjugated OVA peptide. We found that PGP9.5^+^ DRG neurons from naïve female and male mice expressed surface-MHCII (**Fig. 3C**), which co-localized with the OVA-FITC signal (**Fig. 3C**), suggesting OVA peptide is bound to surface-MHCII. Of note, some naïve DRG neurons have diffuse OVA-MHCII signal (**Fig. 3C, solid white arrow)** while others have punctate (polarized) OVA-MHCII staining (**Fig. 3C, dashed white arrow)**. PTX treatment in vivo significantly increased surface-MHCII on cultured neurons compared to neurons from naïve female mice (64.87 ± 2.29 AFUs naïve to 87.59 ± 4.07 AFUs day 14, p=0.0083, n=3) (**Fig. 3D**). In contrast, PTX treatment did not change surface-MHCII on cultured neurons from male mice (59.72 ± 4.22 AFUs naïve to 66.35 ± 0.42 AFUs day 14, p=0.8626, n=3) (**Fig. 3D**). The intensity of OVA increased >2-fold on neurons from day 14 PTX-treated female mice compared to neurons from naïve female mice (41.30 ± 3.10 AFUs naïve to 85.62 ± 4.80 AFUs day 14, p=0.0015, n=3) (**Fig. 3E**). Like surface-MHCII, PTX did not change the intensity of OVA in neurons from male mice (62.77 ± 13.53 AFUs naïve to 77.23 ± 9.51 AFUs day 14, p=0.8563, n=3) (**Fig. 3E**). In addition, the area of polarized MHCII per neuron increased >4-fold after PTX in females (5.5% ± 1.26 naïve to 23.67% ± 1.12 day 14, p=0.0006, n=3) and 2-fold in males (7.3% ± 1.96 naïve to 14.60% ± 0.52 day 14, p=0.0191, n=3) (**Fig. 3F**). Collectively, PTX increases surface-MHCII levels in females and polarization of peptide-bound MHCII in female and male mice, suggesting PTX-induced inflammation increases the likelihood that DRG neurons could activate CD4^+^ T cells particularly in female mice.

### MHCII protein is found in sensory neurons in mouse DRG tissue

MHCII protein was detected in mouse DRG neurons and immune cells **(Fig. S6)** through immunohistochemistry (IHC) visualized by confocal (**Fig. 4A**) and widefield epifluorescence (**Fig. 4B**) microscopy. Confocal microscopy demonstrated that the MHCII signal was detected through the entire DRG neuron cell body (**Fig. 4A**), excluding the possibility that the signal was originating from a proximal non-neuronal cell. DRG neurons from both naïve female and male mice expressed MHCII protein (30.84% ± 6.37 in naïve female DRG and 46.64% ± 10.60 in naïve male DRG) (**Fig. 4B, C**). While PTX and inflammation induce the upregulation of MHCII on immune cells (Pfannenstiel et al., 2010), the effect on neuronal MHCII is unknown. PTX increased the percent of MHCII^+^ DRG neurons in female mice from 30.84% ± 6.37 in naïve to 58.74% ± 7.69 14 days post-PTX, p=0.0190, n=8 (**Fig. 4B, C**). In contrast, the percent of MHCII^+^ neurons in male DRG did not increase significantly after administration of PTX (46.64% ± 10.60 in naïve to 57.98% ± 7.01 14 days post-PTX) (**Fig. 4B, C**).

**Fig.4.**
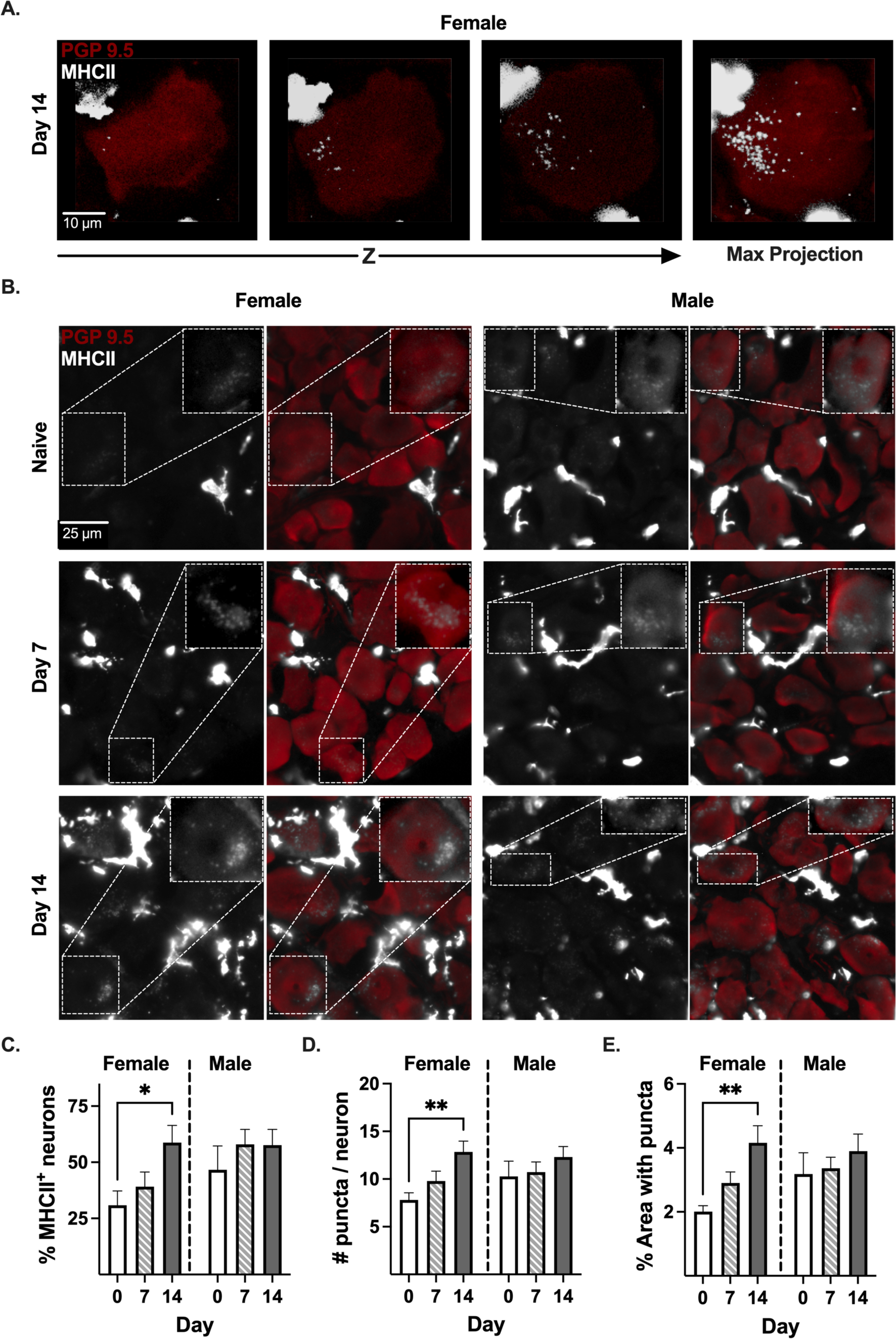
PTX increases neuronal MHCII in female DRG. **(A)** Representative confocal Z-stack and maximum projection IHC images of MHCII (grayscale) in PGP9.5^+^ (red) neuron in L4 DRG tissue from a day 14 PTX-treated female mouse. **(B)** Representative widefield epifluorescence images of DRG tissue from naïve and PTX-treated (day 7 and 14) female and male mice. Inset: representative MHCII^+^ neuron. Quantification of the **(C)** percent of MHCII^+^ neurons, **(D)** number of MHCII puncta/neuron, and **(E)** percent area with MHCII puncta from **(B)**. Statistical significance determined by 2-way ANOVA with Dunnett’s multiple comparison test (*p <0.05, **p < 0.01, n=8/sex).

We observed a distinct punctate staining for MHCII in DRG neurons, a pattern consistent with previous reports (Bosch et al., 2013). Therefore, we quantified the number of MHCII puncta in DRG neurons from naïve and PTX-treated female and male mice. There were 7.61 ± 0.64 MHCII puncta/neuron in the DRG from naïve female mice, which significantly increased to 12.84 ± 1.14 MHCII puncta/neuron 14 days post-PTX (**Fig. 4D,** p=0.0041, n=8). In contrast, the number of MHCII puncta in DRG neurons from male mice remained constant after treatment with PTX (10.28 ± 1.60 in naïve to 12.32 ± 1.1 14 days post-PTX) (**Fig. 4D**). In addition to the number of MHCII puncta, we measured the percent of neuronal cell area with MHCII puncta. The area with MHCII puncta on DRG neurons increased >2-fold in female mice 14 days post-PTX (2.01% ± 0.18 in naïve to 4.16% ± 0.54 14 days post-PTX, p=0.0024, n=8) (**Fig. 4E**). In contrast, the area with MHCII puncta remained constant in DRG neurons from male mice (3.18% ± 0.67 in naïve to 3.90% ± 0.54 14 days post-PTX) (**Fig. 4D**). Collectively, our data demonstrate that PTX treatment increases the expression of MHCII protein in DRG neurons only in female mice.

### PTX induces the expression of MHCII protein in small diameter DRG neurons in female mice

PTX has been shown to affect both large (Aβ) (Zhang and Dougherty, 2014) and small (Aδ and C) (Zhang and Dougherty, 2014) sensory nerve fibers at the peripheral terminals (Siau et al., 2006) and within the DRG (Zhang and Dougherty, 2014, Li et al., 2017). Although low levels of MHCII transcript have been detected across all neuronal subsets in the DRG (Usoskin et al., 2015), it is unknown if the level of MHCII protein is dependent upon the diameter of the neuronal cell body. The diameter of neurons was calculated using an automated analysis pipeline and validated using DRG tissue from TRPV1-lineage td-tomato reporter mice, which identifies TRPV1-lineage neurons as putative nociceptors **(Fig. S7)**. TRPV1 is a broad embryonic marker for many small diameter nociceptors including transient receptor potential cation channel subfamily V member (TRPV1), isolectin-B4 (IB4), and a subset of Aδ neurons (Patil et al., 2018), so all of these nociceptors will express td-tomato protein even though some subpopulations may lose TRPV1 expression after development (i.e., IB4 and Aδ) (Goswami et al., 2014). Although MHCII protein was detected in large diameter neurons (>30µM (Ho et al., 2012)) in female naïve DRG tissue, the majority of MHCII^+^ neurons were small diameter (<25µM: 63.63% ± 3.056) (**Fig. 5A**). After PTX treatment (7 and 14 days), the histogram of neuron diameter size of MHCII^+^ neurons for female mice shifted left toward the diameter mode of td-tomato^+^ neurons from TRPV1-lineage reporter mice (**Fig. 5A**), indicating that PTX induced MHCII protein in small diameter neurons or reduced MHCII in large diameter neurons. The percent of small diameter neurons that were MHCII^+^ increased >2-fold 14 days after PTX treatment (26.89% ± 5.48 in naïve to 55.37% ± 7.72 14 days post-PTX, p=0.0129, n=8**) (Fig. 5B**), while the percent of large diameter neurons that were MHCII^+^ did not increase significantly (43.11% ± 9.23 in naïve to 67.81% ± 7.70 14 days post-PTX**) (Fig. S8)**, indicating that PTX treatment induces MHCII expression in small diameter neurons in female DRG. Moreover, PTX treatment increased the number of MHCII puncta in small diameter neurons from 5.09 ± 0.45 puncta/neuron in naïve female DRG to 8.89 ± 0.87 puncta/neuron 14 days post-PTX (p=0.0022, n=8**) (Fig. 5C**) and increased the percent of neuronal area with MHCII puncta >2-fold (1.98% ± 0.21 in naïve to 4.08% ± 0.55 14 days post-PTX, p=0.0016, n=8**) (Fig. 5D**), further supporting induction of MHCII in small diameter neurons in female DRG after treatment with PTX.

**Fig. 5.**
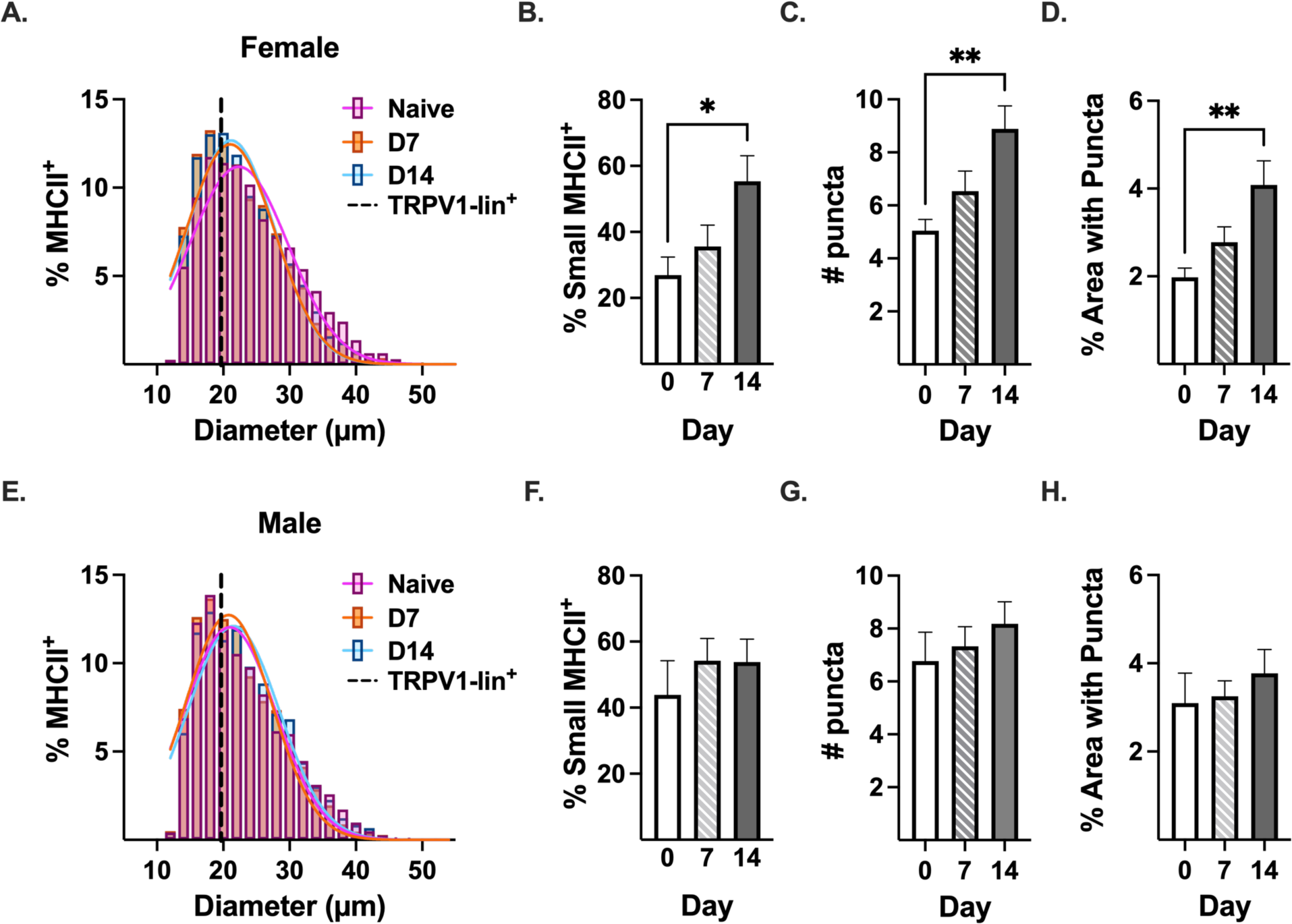
MHCII^+^ DRG neurons are primarily small diameter and expression increases after PTX. **(A, E)** Gaussian distribution of the diameter of MHCII^+^ DRG neurons in DRG tissue from naïve (pink), day 7 (orange) and day 14 PTX-treated (blue) **(A)** female and **(E)** male mice (n=8/sex, pooled neurons). Dashed black line: diameter mode of TRPV1-lineage td-tomato^+^ neurons **(Fig. S7)**. Quantification of **(B, F)** percent of small diameter (≤25 µm) MHCII^+^ DRG neurons positive for MHCII, **(C, G)** number of MHCII puncta/small diameter neuron, and **(D, H)** percent of small diameter neuron area with MHCII puncta. Statistical significance determined by 1-way ANOVA with Dunnett’s multiple comparison test (*p <0.05, **p < 0.01, n=8/sex).

Like female mice, the majority of MHCII^+^ neurons in naïve male DRG were small diameter (<25µM: 66.00% ± 2.02) (**Fig. 5E**). However, unlike females, the administration of PTX did not change the size distribution of MHCII^+^ DRG neurons (pink naïve line overlaps with orange day 7 and blue day 14 PTX lines) (**Fig. 5E**). Therefore, not surprisingly, administration of PTX in male mice did not significantly change the percent of small (43.86% ± 10.38 in naïve to 54.82% ± 6.90 14 days post-PTX**) (Fig. 5F**) or large (55.27% ± 11.06 in naïve to 69.09% ± 7.41 14 days post-PTX**) (Fig. S8)** diameter neurons that were MHCII^+^. Moreover, PTX treatment in male mice did not change the number of puncta/neuron (**Fig. 5G**) or the percent of neuronal area with MHCII puncta (**Fig. 5H**).

### Deficiency in neuronal MHCII prevents the PTX-induced increase in surface MHCII on DRG neurons from female mice

Given the contribution of small nociceptive neurons to CIPN (Li et al., 2017), and that the majority of MHCII is expressed in small diameter neurons, we knocked out one copy of MHCII in TRPV1-lineage neurons (putative nociceptors that include TRPV1, IB4, and a subset of Aẟ neurons; **cHET:** TRPV1^lin^ MHCII^+/-^. As stated previously (**Fig. 3B, 6A)**, 9.66% ± 0.64 of dissociated DRG neurons from naïve WT female mice expressed surface-MHCII. PTX not only induced neuronal surface-MHCII at day 14 (15.44% ± 0.84, p<0.0001, n=9**) (Fig. 3B, 6A)** but also at day 7 (15.07% ± 1.05, p=0.0002, n=7) (**Fig. 6A**). Although DRG neurons from naïve cHET female mice had higher levels of surface-MHCII compared to naïve WT female mice (9.66% ± 0.64 for WT and 18.07% ± 1.44 for cHET, p<0.0001, n=6-9), administration of PTX decreased surface-MHCII in cHET female mice 7 days post-PTX (18.07% ± 1.44 for naïve cHET to 12.57% ± 0.53 for 7 days post-PTX, p=0.0009, n=6**) (Fig. 6A**). By day 14, the frequency of DRG neurons expressing surface-MHCII from cHET female mice returned to baseline levels (12.57% ± 0.53 at 7 days post-PTX to 17.20% ± 0.75 at 14 days post-PTX, p=0.0053, n=6) (**Fig. 6A**). In contrast to female mice, administration of PTX did not change the frequency of surface-MHCII on DRG neurons from WT or cHET male mice (**Fig. 6B**).

**Fig. 6.**
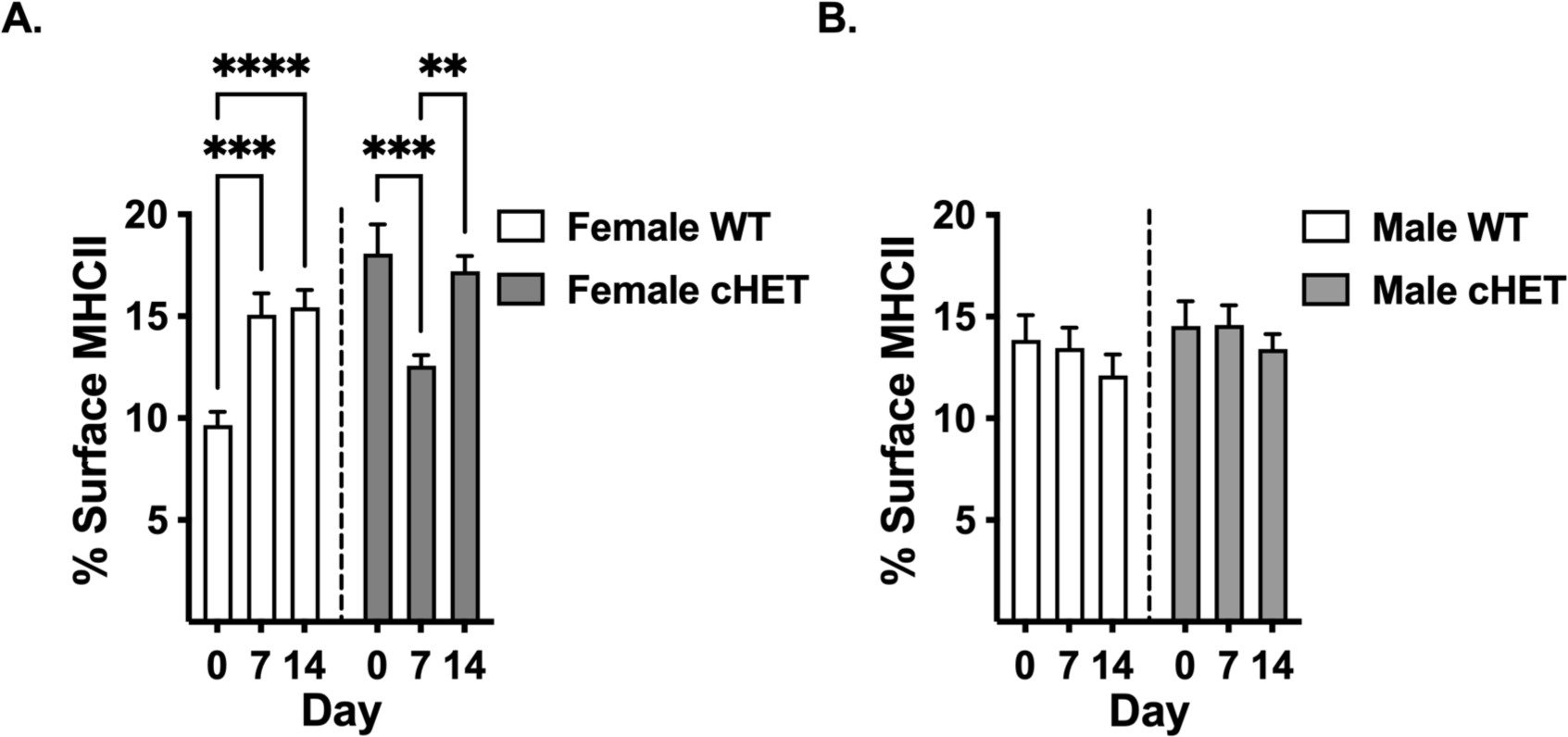
PTX increases surface-MHCII on DRG neurons in WT but not cHET female mice. Flow cytometric analysis of live, dissociated DRG neurons stained with an antibody against MHCII. The frequency of DRG neurons expressing surface-MHCII from wild type and MHCII^+/-^ (cHET) **(A)** female and **(B)** male mice. Statistical significance determined by 2-way ANOVA with Tukey’s multiple comparison test (**p < 0.01, ***p =0.001, ****p <0.0001, n=6-9/sex).

### Both nociceptive and non-nociceptive small diameter neurons express MHCII

Knocking out MHCII in TRPV1-lineage neurons did not change the percent of small diameter MHCII^+^ neurons in the DRG of naïve and day 7 PTX-treated female mice (**Fig. 7A, C**). However, knocking out MHCII in TRPV1-lineage neurons prevented the PTX-induced increase in small diameter DRG neurons in female mice 14 days post-PTX (**Fig. 7B, C**). Knocking out one copy of MHCII in TRPV1-lineage neurons in female mice reduced the percent of small diameter MHCII^+^ neurons by 40% 14 days post-PTX (55.37% ± 7.72 at day 14 in MHCII^+/+^ to 33.30% ± 3.61 at day 14 in cHET, p=0.0231, n=8), and knocking out both copies of MHCII in TRPV1-lineage neurons (**cKO:** TRPV1^lin^ MHCII^-/-^) reduced the percent of small diameter MHCII^+^ neurons by 68% 14 days post-PTX (55.37% ± 7.72 at day 14 in MHCII^+/+^ to 17.99% ± 3.81 at day 14 in cKO, p=0.0015, n=8 WT and n=4 cKO) (**Fig. 7C**), further supporting that PTX induces MHCII in small diameter nociceptive DRG neurons. However, as TRPV1-lineage neurons comprise only 65% of small diameter neurons **(Fig. S7C)**, this indicates that the residual MHCII in small diameter neurons of cKO mice is a result of expression in a population of small non-nociceptive neurons in the DRG of both naïve and PTX-treated female mice.

**Fig. 7.**
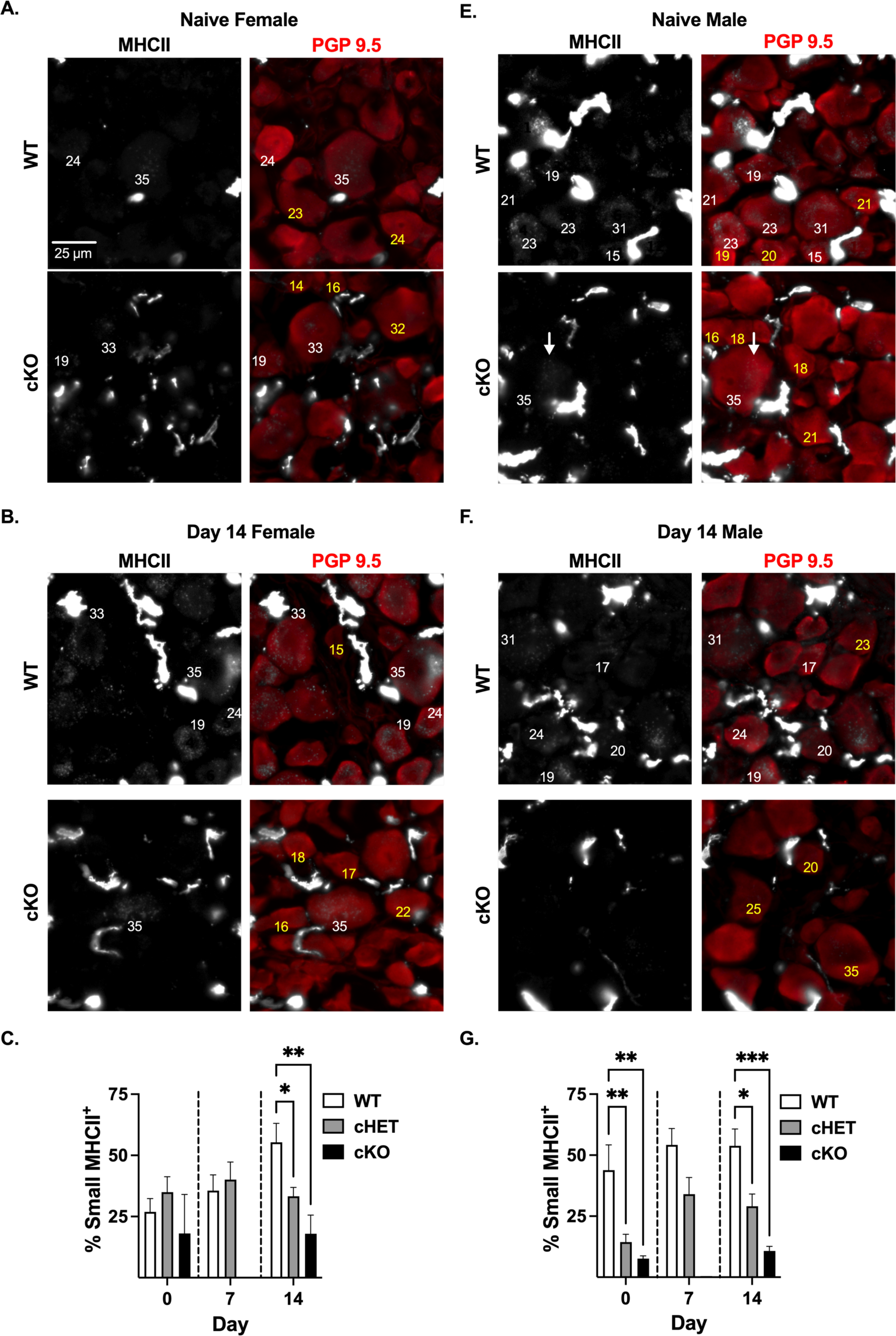
TRPV1-lineage neurons express MHCII in naïve male DRG, and PTX treatment induces MHCII in TRPV1-lineage neurons in female DRG. **(A-F)** Immunohistochemistry of MHCII (grayscale) and neuronal marker PGP9.5 (red) in L4 DRG tissue from TRPV1-lineage MHCII^+/+^ (WT), MHCII^+/-^ (cHET) and MHCII^-/-^ (cKO) naïve and day 14 PTX-treated female and male mice. Number = neuron diameter size, yellow (MHCII^-^) and white (MHCII^+^). Percent of small diameter MHCII^+^ neurons for **(C)** female and **(G)** male WT, cHET, and cKO mice. Statistical significance determined by 2-way ANOVA with Dunnett’s multiple comparison test (*p <0.05, **p <0.01; n=7-8 for WT and cHET; n=4 for naïve and day 14 cKO, n=0 for male and female day 7 cKO).

In contrast to female mice, knocking out MHCII^+^ in TRPV1-lineage neurons in male mice reduced the percent of small diameter MHCII^+^ neurons by 66% in naïve cHET mice (43.86% ± 10.38 in MHCII^+/+^ to 14.38% ± 3.18 in cHET, p=0.0039, n=8) and 82% in naïve cKO mice (43.86% ± 10.38 in MHCII^+/+^ to 7.62% ± 1.16 in cKO, p=0.0038, n=8 WT and n=4 cKO) (**Fig. 7E, G**). Likewise, knocking out one copy of MHCII^+^ in TRPV1-lineage neurons in male mice reduced the percent of small diameter MHCII^+^ neurons by 36% at day 7 (54.19% ± 6.75 at day 7 in MHCII^+/+^ to 34.04% ± 6.80 at day 7 in cHET, p=0.0164, n=8) and 45% at day 14 (53.82% ± 6.90 at day 14 in MHCII^+/+^ to 29.09% ± 5.03 at day 14 in cHET, p=0.0164, n=8 WT). Furthermore, knocking out both copies of MHCII in TRPV1-lineage neurons in males reduced the percent of small diameter MHCII^+^ neurons by 80% 14 days post-PTX (53.82% ± 6.90 at day 14 in MHCII^+/+^ to 10.77% ± 1.85 at day 14 in cKO, p-0.0006, n=4 cKO) (**Fig. 7E, G**). Reduction of MHCII in TRPV1-lineage neurons (TRPV1, IB4, and a subset of Aẟ neurons) further supports novel MHCII expression in DRG neurons from both sexes but regulation by PTX in only female mice.

### MHCII-expressing nociceptors attenuate cold hypersensitivity

Given that small diameter nociceptors contribute to CIPN (Li et al., 2017) and cold hypersensitivity is one of the major symptoms (Luo et al., 2019, Polomano et al., 2001, Smith et al., 2004), we determined the extent MHCII expressed in small nociceptive neurons protects against cold hypersensitivity by the thermal placed preference (TPP) behavioral test. Although the mice are motivated to explore their environment, mice that are hypersensitive will spend progressively less time on the cold plate with each successive trial as a result of learning. A faster acquisition of a learned avoidance response demonstrates a greater degree of cold hypersensitivity.

Naïve female mice were slightly hypersensitive to the cold plate only at trial 2 (53.90% ± 2.16 at baseline (BL) to 36.14% ± 4.24 at trial 2, p=0.0075, n=13) while knocking out one copy of MHCII in TRPV1-lineage neurons (cHET) induced a slight increase in cold hypersensitivity at trial 4 (46.87% ± 2.86 at BL to 27.71% ± 6.92 at trial 4, p=0.0127, n=16) (**Fig. 8A**). In contrast, reducing MHCII expression in TRPV1-lineage neurons (cHET) drastically increased cold hypersensitivity in naïve male mice (**Fig. 8B**), which is consistent with MHCII being primarily expressed in small nociceptive neurons in naïve male mice (**Fig. 7G**). With each successive trial, the TRPV1-lineage cHET male mice spent less time on the cold plate with a significant difference between BL (49.52% ± 3.60) and trial 3 (20.67% ± 5.33, p=0.0024, n=12) but was not significantly different from wildtype (WT) mice until trial 4 (36.55% ± 7.35 for WT and 11.49% ± 5.00 for cHET, p=0.0377, n=12-15) (**Fig. 8B**). This enhanced learning pattern to avoid the cold plate in cHET male mice indicates cold hypersensitivity, demonstrating that expression of MHCII in small nociceptive neurons prevents cold hypersensitivity in only naïve male mice.

**Fig. 8.**
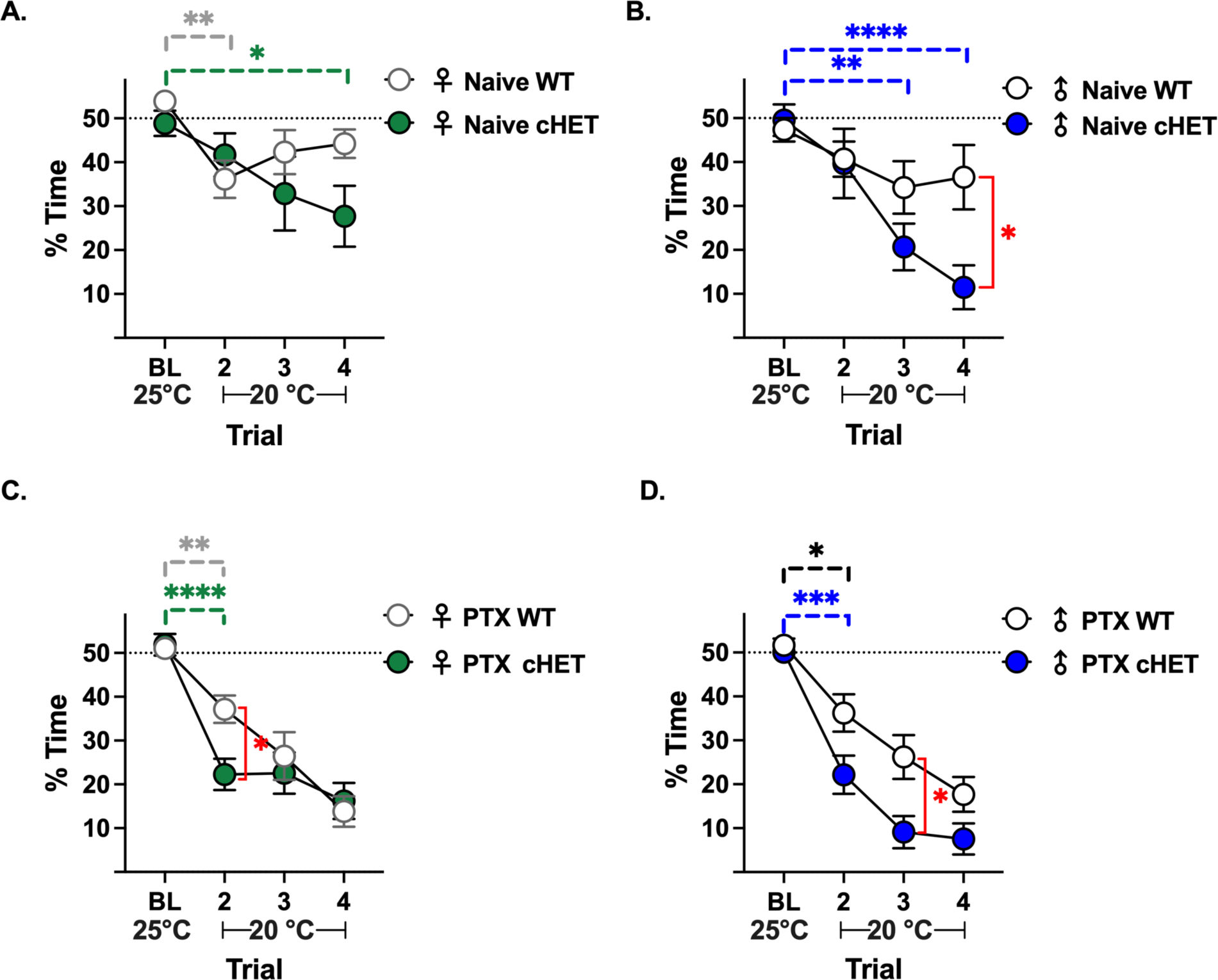
MHCII in TRPV1-lineage neurons attenuates cold hypersensitivity. Thermal placed preference test (test plate = 25 °C, baseline (BL) and 20 °C, reference trials 2-4): Percent of time spent on test plate for **(A, B)** naïve and 6 days post-PTX **(C, D)** WT and cHET female and male mice. Dashed line: expected percent of time on test plate when mice are not hypersensitive to cold. Statistical significance determined by repeated measures mixed-effects model (REML) with Dunnett’s multiple comparison test comparing trials 2-4 to BL within same group (horizontal bars, complete statical analysis in **Table S1**), and Sidak’s multiple tests for significance between groups at the same trial (red vertical bars) (*p <0.05, **p <0.01; ****p<0.0001 n=12-15/sex/group).

Administration of PTX induced cold hypersensitivity in WT (51.04% ± 1.77 at BL to 37.16% ± 3.11 at trial 2, p=0.0072, n=12) and cHET (51.87% ± 2.45 at BL to 22.28% ± 3.58 at trial 2, p<0.0001, n=12) female mice by trial 2; however cHET female mice were statistically more hypersensitive than WT female mice (37.16% ± 3.11 for WT and 22.28% ± 3.58 for cHET, p=0.0193, n=12) (**Fig. 8C**). Like female mice, PTX-treated cHET male mice were more hypersensitive to cold temperature than PTX-treated WT male mice. Administration of PTX induced cold hypersensitivity in WT (51.66 ± 1.53 at BL to 36.23 ± 4.27, p=0.0125, n=15) and cHET (50.10 ± 2.21 at BL to 22.17 ± 4.36 at trial 2 in cHET, p=0.0004, n=12) male mice at trial 2 (**Fig. 8D**); however, cHET male mice were only more hypersensitive to cold temperature compared to WT male mice at trial 3 (26.22% ± 4.99 for WT and 9.11% ± 3.67 for cHET, p=0.0424, n=12-15) (**Fig. 8D**). Novel expression of MHCII in small diameter TRPV1-lineage nociceptors (TRPV1, IB4, and a subset of Aẟ) attenuates PTX-induced cold hypersensitivity in both female and male mice.

## DISCUSSION

The debilitating nature of CIPN has been a major obstacle for the continuation of life-saving measures for individuals with cancer. Recently, it has been shown that anti-inflammatory cytokines (IL-10 and IL-4) are protective against CIPN (Shi et al., 2018, Krukowski et al., 2016); however, systemic use of IL-10 in general has had poor clinical outcomes (Saxena et al., 2015) as it can promote the progression of cancer (Lech-Maranda et al., 2006) and propagate chronic viral infections (Nelson et al., 2003). In our recent work, we found that PTX induces a robust increase in anti-inflammatory CD4^+^ T cells in the DRG of female mice (Goode et al., 2022), but the mechanism by which CD4^+^ T cells are activated and the extent cytokines released by CD4^+^ T cells target neurons in the DRG are unknown. We hypothesize that PTX exposure increases MHCII on sensory neurons, which would stimulate the paracrine release of anti-nociceptive cytokines by CD4^+^ T cells to suppress CIPN.

MHCII is traditionally thought to be constitutively expressed only in APCs but can be induced by inflammation in non-APCs (van Velzen et al., 2009). RNA seq data sets (Nguyen et al., 2021, Tavares-Ferreira et al., 2022, Usoskin et al., 2015, Lopes et al., 2017) demonstrate that mouse and human DRG neurons express transcripts for MHCII and MHCII-associated genes. However, there are no reports to date that demonstrate MHCII protein expression in terminally differentiated neurons. Neuronal MHCII was most likely overlooked in the P. Hu et al. study as MHCII^+^ cells in the DRG were identified based on morphology instead of co-staining with cell-specific markers. Additionally, the signal intensity of immune cell MHCII is >5 times greater than neuronal MHCII, indicating a lower density of MHCII molecules in neurons. Therefore, if not actively investigating neuronal MHCII, the settings used to capture immune cell MHCII would obscure neuronal MHCII. In addition, trying to identify a few molecules of MHCII is technically challenging as the signal is not much higher than background. Thus, it is quite possible that neurons with extremely low level of MHCII were classified as negative cells in our study. Even though DRG neurons have fewer MHCII molecules per cell area than non-neuronal cells, it is likely still functionally significant given that clustering allows for as little as 210 MHCII molecules to activate a CD4^+^ T cell (Banchereau and Steinman, 1998, Harding and Unanue, 1990). Novel expression of MHCII in neurons will provide insight into the mechanism by which CD4^+^ T cells contribute to pain, autoimmunity, and neurological diseases.

DRG neurons express RNA transcripts for chemokines CCL19/21/27 (Lopes et al., 2017), which are potent chemoattractants for CD4^+^ T cells. Recently, CCL19/21 have been shown to induce the migration of CD4^+^ T cells to non-lymphoid tissue, especially to the nervous system (Kivisäkk et al., 2003, Axtell and Steinman, 2009), to preserve the balance between immune surveillance (against tumor and pathogens) and tolerance. The presence of CD4^+^ T cells in naïve mouse DRG (Goode et al., 2022, Liu et al., 2014), together with the expression of MHCII in neurons support direct cell-cell interaction, despite SGCs surrounding DRG neurons (Dixon, 1969). Our present work demonstrates that CD4^+^ T cells can breach the SGC barrier, most likely through a natural gap in the surrounding glial envelope or a gap junction between adjacent glial cells. CD4^+^ T cells in close proximity to DRG neurons in the naïve mouse suggests a role for CD4^+^ T cells in not only immune surveillance and tolerance, but also regulating neuronal function.

Nerve injury and nerve transection models demonstrate that macrophages can breach SGCs and lie directly against the neuron in the DRG (Hu and McLachlan, 2002) and trigeminal ganglion (Iwai et al., 2021). Similarly, we found CD4^+^ T cells breaching the SGCs barrier after treatment with PTX. Furthermore, we observed that CD4^+^ T cells appeared flattened on the side closest to the neuron (**Fig. 1B**, **C**, **S2, S3**), which resembles the CD4^+^ T cell shape that occurs during the formation of the immunological synapse (IS) with an APC (Delon et al., 1998, Donnadieu et al., 1994, Lin et al., 2015). TCR signaling molecules accumulate at the flattened contact site resulting in an increase in calcium signaling, which is required to activate the T cell (Lin et al., 2015). Future studies are needed to determine whether DRG neurons form an IS with CD4^+^ T cells to induce TCR signaling. Neuronal MHCII-dependent CD4^+^ T cell activation would target cytokine release toward neurons to suppress hypersensitivity after inflammation and/or nerve injury.

There are known sex difference in immune and nociceptive responses. For example, although DRG neurons express transcripts and/or protein for TLRs 1-9 (Lopes et al., 2017, Wang et al., 2020, Cameron et al., 2007, Barajon et al., 2009, Xu et al., 2015, Zhang et al., 2018), female mice have >5-fold more TLR4-6 transcripts than male mice (Lopes et al., 2017). Further supporting a sex-specific role for TLR4 in the DRG, Thomas A Szabo-Pardi et al demonstrate that TLR4 signaling in DRG neurons mediates mechanical hypersensitivity after a nerve injury in only female mice (Szabo-Pardi et al., 2021). Here we report a sex-specific difference in neuronal MHCII, though the mechanism of MHCII induction in only females after PTX treatment remains unknown. TLR4 binds to PTX (Byrd-Leifer et al., 2001) and has been shown to regulate MHCII in APCs (Rathinam et al., 2009), together suggesting one potential mechanism by which MHCII is induced in DRG neurons in female mice.

Although the major role for surface MHCII is to activate CD4^+^ T cells, cAMP/PKC signaling occurs in the MHCII-expressing cell(Harton, 2019). In addition, it has recently been shown that endosomal MHCII plays an important role in promoting TLR responses(Liu et al., 2011), and since DRG neurons are known to express TLRs (Lopes et al., 2017, Wang et al., 2020, Cameron et al., 2007, Barajon et al., 2009, Xu et al., 2015, Zhang et al., 2018), this suggests the potential for T-independent MHCII^+^ neuron responses. Knocking out one copy of MHCII in TRPV1-lineage neurons (cHET) from female mice did not change total MHCII 7 days post-PTX but reduced surface-MHCII. Accordingly, PTX-treated cHET female mice were more hypersensitive to cold than cHET naïve and WT female mice, suggesting a role for neuronal MHCII in CD4^+^ T cell activation and/or neuronal cAMP/PKC signaling. In contrast, knocking out one copy of MHCII in TRPV1-lineage neurons (cHET) from male mice did not change surface-MHCII in naïve or PTX-treated mice but reduced total MHCII, indicating endosomal MHCII and potentially a role in TLR signaling. Future studies are required to delineate MHCII surface and endosomal signaling mechanisms in naïve and PTX-treated female and male mice.

Small diameter neurons, which consist of Aẟ and unmyelinated C-fibers, have also been shown to contribute to CIPN (Li et al., 2017). C-fibers can be further subdivided into nociceptive TRPV1 and IB4 neurons, which respond to noxious stimuli, and non-nociceptive **l**ow **t**hreshold **<u>m</u>**echano**<u>r</u>**eceptors (C-LTMRs: majority tyrosine hydroxylase (TH) neurons and minority mas-related G-protein receptor B4 (MrgB4) neurons). As cold hypersensitivity is a major symptom of CIPN (Luo et al., 2019, Polomano et al., 2001, Smith et al., 2004), we assessed the contribution of MHCII in small diameter nociceptive neurons by TPP. TPP is a behavioral test that incorporates the ability to choose and the potential to learn, demonstrating higher order pain processing. We found that reducing the expression of MHCII in TRPV1-lineage neurons in naive male and female mice induced cold hypersensitivity in only male mice. Therefore, the majority of MHCII^+^ neurons in the DRG of naïve female mice were not TRPV1-lineage neurons but non-nociceptive C-LTMRs. Although C-LTMRs typically process innocuous gentle and affective touch, there are a few reports that indicate a role for TH neurons in pain (Malin et al., 2007, François et al., 2015), suggesting that either MHCII in TH neurons protects against cold hypersensitivity or that neuronal MHCII does not contribute to cold hypersensitivity in naïve female mice.

Reducing the expression of MHCII in TRPV1-lineage neurons exacerbated PTX-induced cold hypersensitivity in both male and female mice. Future studies will evaluate the role of MHCII in PTX-induced mechanical hypersensitivity, another prominent feature of CIPN. In addition to TRPV1-lineage neurons, we found in both female and male mice that MHCII is expressed in large diameter neurons, which primarily convey non-nociceptive input (proprioception, touch, pressure, vibration) from the periphery to the spinal cord but may contribute to nociceptive circuits following tissue or nerve injury as in CIPN (Zhang and Dougherty, 2014). MHCII expression in small and large LTMRs indicates that PTX-induced cold hypersensitivity may be even greater if MHCII is eliminated from all neurons and suggests a role in proprioception, pressure, and touch (light, gentle and affective). Future studies will address the role of MHCII in large and small diameter LTMRs in pain and other sensory modalities.

Neuronal MHCII represents a novel mechanism to suppress CIPN, which could be exploited for therapeutic intervention against not only pain but autoimmunity and neurological diseases. Moreover, future investigation will evaluate the extent neuronal MHCII facilitates communication with CD4^+^ T cells and MHCII T-independent signaling in DRG neurons during homeostasis and inflammation/nerve injury.

## MATERIALS AND METHODS

### Mice

Male and female mice were purchased from Jackson Laboratory **(Table S2)**. Male TRPV1^Cre^ mice were bred to female MHCII^fl/fl^ mice to generate TRPV1^lin^ MHCII^+/-^ heterozygote mice **(cHET)**. cHET female mice were mated back to MHCII^fl/fl^ male mice to delete MHCII from TPRV1-lineage neurons, which identifies putative nociceptors that include IB4, TRPV1, and a subset of Aẟ neurons (TRPV1^lin^ MHCII^-/-^ mice; **cKO**). To confirm the presence of MHCII^fl/fl^ and Cre, mouse tails were digested with 50mM NaOH/50mM HCl/1M Tris-HCl, pH 8.0. Primers (Table S3) and Promega GoTaq® G2 Hot Start Polymerase were used in PCR to detect MHCII^fl/fl^ (199 base pair), MHCII^fl/+^ (199 and 295 base pairs), and wild type MHCII^+/+^ (295 base pair) mice. PCR using GoTaq® Polymerase determined the presence of Cre **(Table S3)**. cHET×MHCII^fl/fl^ crosses only yielded 8% cKO mice (4% per sex) instead of the predicted 25% (12.5% per sex) based on normal Mendelian genetics. Thus, cKO mice were only used to validate MHCII protein in small nociceptive neurons.

All mice were given food and water *ad libitum*. All experimental protocols followed National Institutes of Health guidelines and were approved by the University of New England Institutional Animal Use and Care Committee. Mice were euthanized with an overdose of avertin (0.5cc of 20 mg/ml) followed by transcardiac perfusion with 1× phosphate buffered saline (PBS). This method of euthanasia is consistent with American Veterinary Medical Association Guidelines for the Euthanasia of Animals.

### Paclitaxel and Thermal Place Preference test (TPP)

PTX (Sigma-Aldrich) was solubilized in cremophor:ethanol 1:1 and diluted 1/3 in 1× PBS. A single injection of 6mg/kg PTX at a volume of 10ml/kg bodyweight was given on day 0 as done previously (Liu et al., 2014, Goode et al., 2022). Mice were randomly assigned a treatment, which experimenters were blinded to during testing. The TPP test was used to quantify non-evoked measures of cold hypersensitivity before and after PTX. Mice were habituated to the room and testing apparatus, which includes two adjustable thermal plates (test and reference). Mice were placed on the reference plate to begin each 3-minute trial. The reference plate side was the same for all trials for an individual mouse, but the sides for the reference and test plates were counterbalanced within each group to exclude a potential side preference. For the habituation trial (#1), both the test and reference plates were set to 25 °C, and the mice were allowed to explore for 3 minutes. Any mouse that spent <30 seconds on the test plate during the habituation trial was excluded from further testing. For trials #2-4, the reference plate was set to 25 °C and the test plate to 20 °C. The percent of time the mice spent on the test plate was reported for each trial. Separate cohorts of mice were run for naive and 6 days post-PTX to prevent confounding effects of avoidance learning during the naive session impacting the day 6 results.

### Immunohistochemistry (IHC)

Mice were perfused with 1× PBS followed by 4% paraformaldehyde (PFA; Sigma-Aldrich). L4 DRGs were post-fixed in 4% PFA for 1-2 hours, then transferred to 30% sucrose/0.02% sodium azide at 4 °C. DRGs were embedded in Clear Frozen Selection Compound prior to serially sectioning at 12 µm slices. DRGs were permeabilized with 1× PBS/0.1% Triton X-100 (PBS-T; Sigma-Aldrich) for 15 minutes at room temperature (RT) and incubated with blocking buffer (PBS-T plus 5% normal donkey serum) for 1 hour at RT.

Slides were incubated with primary antibodies **(Table S4)** in blocking buffer overnight at RT in a humidified, light-protected chamber. Slides were washed 3X with 1× PBS and secondary antibodies **(Table S4)** added for 1 hour at RT. Slides were washed 3X with 1× PBS, then cover slipped with Fluoroshield Mounting Medium (±DAPI, Abcam). RFX1 images were acquired with a 20× objective (HC PL Fluotar NA 0.40) on an ImageXpress Pico (Molecular Devices) automated widefield fluorescence microscope with a high-sensitivity, 5-megapixel CMOS camera and using CellReporterXpress Software (CRX; version 2.9.3). Widefield CD4^+^ T cell–DRG neuron images were acquired with a 40× objective (HC PL Fluotar NA 0.60 with correction ring; Z-stacks with 1 µm step) on an ImageXpress Pico and processed by Best Focus projections in CRX prior to export. MHCII images were acquired with a 40× objective (Plan Fluor NA 0.75) on a Keyence BZ-X710 automated widefield fluorescence microscope with incorporated cooled monochrome CCD camera (960×720 image resolution, 14-bit gradation). Confocal images of CD4^+^ T cell-DRG neuron contacts and neuronal MHCII IHC and neuronal MHCII IHC were acquired with a Leica TCS SP5 II confocal microscope using a 40× objective (HCX PL APO NA 1.3 under oil, RI 1.52, zoom 3.0, 0.34 µm step size, 12-bit, scan format either 512×512 or 1024×512). 3D volume of CD4^+^ T cell-DRG neuron contact was generated by MetaXpress software (Molecular Devices, version 6.7.1.157) using quadratic blending for channel display.

### IHC Automated Image Analysis

IHC analysis was performed using an automated pipeline to eliminate human bias and validated manually by two independent blinded experimenters.

#### MHCII^+^ DRG neurons

Keyence images were imported into Fiji (Schindelin et al., 2012) to generate single channel images, then analyzed by MetaXpress software. The percent of MHCII^+^ DRG neurons was quantified by developing a Custom Module within MetaXpress. MHCII^+^ DRG neurons were identified as follows: 1) A mask was generated to capture neurons within the DRG tissue while excluding nerve fibers. 2) A subsequent mask was created to identify single neurons based on size and PGP9.5 intensity. 3) A “Shrink” function, which decreased the borders of the neuron mask by 4 pixels, was applied to exclude potential MHCII signal from surrounding SGCs. 4) A mask to capture MHCII^+^ non-neuronal cells was created based on size and signal intensity, which was >5 times greater than neuronal MHCII (≥25,000 arbitrary fluorescent units (AFUs)). 5) The “Grow” function, which expanded the borders of the non-neuronal mask by 7 pixels, was applied to capture low-intensity non-neuronal MHCII staining. 6) To exclude non-neuronal MHCII signal that overlapped DRG neurons, a logical operation was applied. Neurons excluding non-neuronal cell overlap were used to calculate MPI of MHCII in each neuron. 7) The threshold for MHCII^+^ DRG neurons was set as the mean intensity for the MHCII signal at the 99^th^ percentile for DRG L4 stained with an isotype control antibody (n = 3 naïve females). Automated analysis allowed for consistent measurement of MHCII expression across all conditions (872.1 ± 25.28 neurons/mouse).

#### MHCII puncta and neuron diameter

MHCII puncta were identified within the neuronal mask as regions of ≥1.5 pixels containing MHCII signal ≥800 MPI above local background. The number of MHCII puncta was determined by measuring the number of puncta per neuron (**Fig. 4D, 5C, 5G).** The percent of neuronal area with MHCII puncta was calculated by dividing the sum of puncta-containing area (pixel^2^) by the total area of the neuron (**Fig. 4E, 5D, 5H**). The diameter of DRG neurons was determined by: (1) Measuring the total area of each neuron (pixel^2^); (2) Multiplying the area (pixel^2^) by 6.718 µm^2^/pixel^2^ to convert to µm^2^; (3) using the equation (A = πr^2^).

#### RFX1^+^ DRG neuron

Images were imported into MetaXpress analysis software for automated analysis using a Custom Module as follows: 1) As described for the neuronal MHCII^+^ automated analysis, masks were created to identify DRG tissue and neurons. 2) Separate masks for non-neuronal and neuronal nuclei were created based on nuclei size and intensity of DAPI. 3) Non-neuronal nuclei were removed from the neuron mask using a logical operation. 4) The number of PGP9.5^+^ neurons with a visible nucleus that had nuclear RFX1 with a MPI >37.19 were counted and divided by the total number of neurons with visible nuclei to determine the percent of RFX1^+^ DRG neurons.

### Immunocytochemistry (ICC)

DRGs collected from naïve and day 14 PTX-treated male and female mice were pooled for each mouse and acutely dissociated into a single cell suspension as described previously (Goode et al., 2022). DRG neurons were isolated by depleting non-neuronal cells through MACS following the manufacturer’s protocol (Miltenyi). MACS-DRG neurons were resuspended in F12 media supplemented with 10% fetal bovine serum (FBS) and 100 µg/ml primocin (Invivogen) and plated on chamber slides (Millipore) coated with 20 µg/ml laminin (Sigma) and 50 µg/ml poly-D-lysine (<u>Thermo</u>).

#### Total MHCII

After an 18-hour incubation at 37 °C, MACS-neurons were washed with 1× PBS and fixed with 4% PFA. Cells were washed with 1× PBS and incubated with blocking buffer for 30 minutes at RT. PGP9.5 was added to the cells for 1 hour at RT. Cells were washed with 1× PBS and incubated with anti-rabbit 488 secondary antibody and MHCII for 1 hour at RT. Slides were imaged with a Leica TCS SP5 II confocal microscope using a 40× objective under oil.

#### Extracellular MHCII and OVA peptide

After an 18-hour incubation at 37 °C, media was replaced with F12/10% FBS/100 µg/ml primocin. FITC-conjugated OVA peptide (Anaspec) was added at 10 µg/ml for 30 minutes at 37 °C. Cells were washed with 1× PBS/1% FBS and blocked with CD16/32 (Biolegend). After 10 minutes, cells were incubated with MHCII for 30 minutes on ice to stain for extracellular MHCII. Cells were washed, fixed with 4% PFA, and stained with PGP9.5. Slides were imaged on the Keyence microscope with a 20× objective.

#### ICC Automated Image Analysis

MHCII on the surface of cultured DRG neurons was quantified using a Custom Module developed in the MetaXpress analysis software. DRG neurons were identified using a mask based on cell size and PGP9.5 intensity. Then the MPI for MHCII and OVA were measured for each neuron. MHCII polarization was quantified as the accumulation of surface MHCII molecules within a defined area (≥ 2 pixels) with a signal intensity ≥ 1.7-fold greater than local background. Polarized MHCII area was divided by the total DRG neuron area to give the percent neuron area containing polarized MHCII.

### Flow cytometry

DRG cells from naïve, day 7, and day 14 PTX-treated female and male WT and cHET mice were incubated at RT for 30 min with Live/Dead Fixable Violet (Thermo). Cells were pre-incubated with CD16/32 for 10 min prior to the addition of MHCII or isotype for 20 min at 4 °C. Cells were washed with FACs buffer (1× PBS, 1% FBS) and fixed at RT for 20 min with Intracellular Staining Fixation Buffer (BioLegend). Cells were washed with 2× Cyto-FastTM Perm Wash solution (CFPWS; BioLegend) and incubated with PGP9.5 for 20 min at RT. Cells were washed with 2× CFPWS and incubated with donkey anti-rabbit Cy3 for 20 min at RT. VersaComp Antibody Capture Kit (Beckman Coulter) was used to set compensation to correct for spectral overlap. Unstained DRG cells were used to check for auto-fluorescence. Beckman Coulter Cyto-FLEX S System B2-R3-V4-Y4 was used to acquire >10,000 events in the live PGP9.5 DRG neuron gate. Data was analyzed with FlowJo^TM^ 10.8.2.

### Western blot

MACS-DRG neurons from naïve and day 14 PTX-treated female and male mice were lysed in DiGE lysis buffer (7 M urea, 2 M thiourea, 4% CHAPS, 30 mM Tris-HCl) supplemented with 1× Halt Protease and Phosphatase Inhibitor Cocktail (Thermo). Lysates were quantified with the EZQ Protein Quantification Kit (Thermo). Lysates (20 μg) were added to Bolt bis-Tris 12 % gel (Thermo). SDS-PAGE was performed using 1× MOPS SDS Running Buffer (Life Technologies). Proteins were transferred to Immobilon-FL PVDF membrane (Millipore) using 1× Power Blotter 1-Step Transfer Buffer (Invitrogen). PVDF membranes were incubated at 4 °C for 48 hours with MHCII and beta tubulin as the loading control, then incubated with 1:2000 donkey anti-rabbit Cy5 secondary antibody for 2 hours at RT. PVDF membranes were imaged using the Typhoon 9600 laser scanner (GE). Band intensities were quantified using AutoQuant imaging software.

### Statistical Analysis

All experiments were analyzed using Graphpad Prism 9 (Graphpad Software, Inc) and reported as mean ± standard error of mean. An arcsine transformation was performed on all percentage data prior to statistical analysis. A two-way ANOVA with Sidak’s multiple comparison test was used to compare the means of neuronal MHCII from naïve and day 14 PTX-treated female and male mice for western blot, ICC, and flow cytometry (**Figs. 2, 3**). An unpaired t-test was used to compare the means of neuronal RFX1 for male and female mice (**Fig. S4**). A two-way ANOVA with Dunnett’s multiple comparison test was used to compare the means of MHCII for PTX-treated (day 7 and 14) male and female mice to the naïve control (**Figs. 4 and 7**). A one-way ANOVA with Dunnett’s multiple comparison test was used within each sex to compare the mean of the percent of small/large diameter MHCII^+^ DRG neurons from PTX-treated mice (day 7 and 14) to the naïve control (**Fig. 5 and S8**). TPP behavior was analyzed using a repeated measures mixed-effects model (REML) with Dunnett’s multiple comparison test comparing the mean percent of time on the cold plate (trials #2-4) to the control (#1) within the same group. Sidak multiple comparison test was used to compare the means between two groups within the same trial (**Fig. 8**).

## Supplementary Materials

### Tables

Table S1. Two-way ANOVA analysis with Dunnett’s multiple comparison test for PTX-treated female and male WT and cHET for TPP test shown in Fig. 7.

Table S2. Jackson Laboratory mouse strains used in the study.

Table S3. Primer sequences for TRPV-1 lineage MHCII conditional knockout mice.

Table S4. Antibody details for each application.

### Figures

Fig. S1. CD4^+^ T cells are found in close proximity to neurons in naïve female DRG tissue.

Fig. S2. CD4^+^ T cell breaches the SGC barrier in female mouse DRG tissue.

Fig. S3. CD4^+^ T cells are in close proximity to neurons in female mouse DRG tissue.

Fig. S4. DRG neurons from naive female and male mice express nuclear RFX1 protein.

Fig. S5. PGP9.5^+^ neurons comprise a fraction of total DRG cells.

Fig. S6. CD11b/c^+^ immune cells in DRG tissue express MHCII.

Fig. S7. High throughput automated image analysis module identifies small and large diameter neurons in female and male DRG tissue.

Fig. S8. PTX does not change the percent of large diameter neurons that express MHCII in male and female DRG.

## Acknowledgments

We would like to thank the UNE COBRE Behavior Core, especially Dr. Tamara King and Denise Giuvelis, for running TPP and assisting in experimental design, analysis, and interpretation.

## Funding

National Institutes of Health grant R01CA267554 (DG)

National Institutes of Health grant P20GM103643 (IM, DG)

University of New England, Office of Sponsored Programs mini grant 17076 (DG)

## Author contributions

Conceptualization: DJG

Methodology: DJG, EEW, NEM

Software: EEW

Investigation: EEW, DJG, NEM

Validation: NEM, RCC

Visualization: DJG, EEW

Funding acquisition: DJG

Project administration: DJG

Supervision: DJG

Writing – original draft: EEW, DJG

Writing – review & editing: EEW, DJG, RCC

## Competing interests

The authors declare no competing interests.

## Data and materials availability

All data are available in the main text or the supplementary materials.

## Notes

### Competing Interest Statement

The authors have declared no competing interest.

### Summary of Updates

Quantification of CD4+ T cell/mm2 DRG tissue was added to Figure 1. Male data were added to Figures 2 and 3. Day 7 data added to histograms in Figure 5 and graphs in Figure 7. New Figure added: Fig. 6. PTX increases surface-MHCII on DRG neurons in WT but not cHET female mice.

