## Supplementary figures and images for "Novel expression of MHC II in DRG neurons attenuates paclitaxel-induced cold hypersensitivity in male and female mice"

### Fig. S2 3D Video

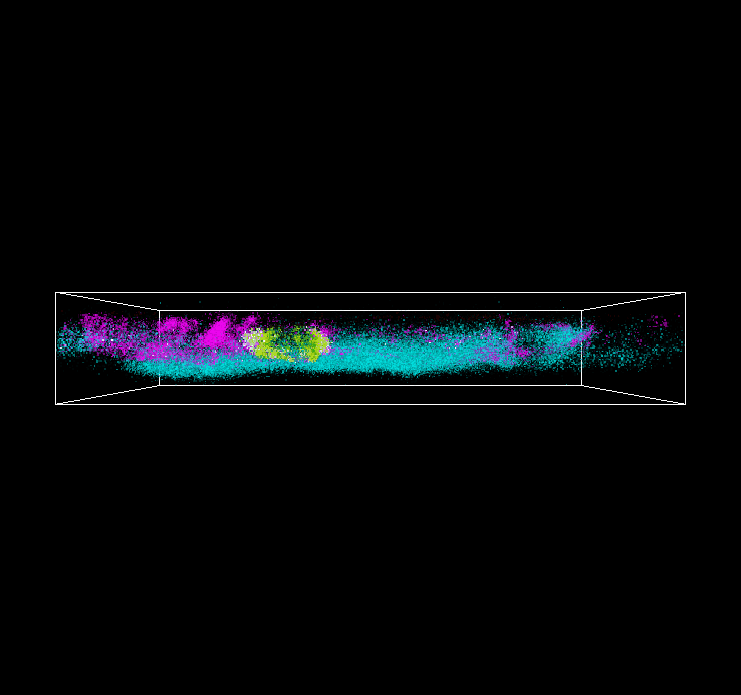
