## Supplementary Materials for "Novel expression of MHC II in DRG neurons attenuates paclitaxel-induced cold hypersensitivity in male and female mice"

### Supplemental Tables

| Group | Dunnett's multiple comparisons test | Summary | P Value |
|---|---|---|---|
| cHET Female PTX | 1 vs. 2 | **** | <0.0001 |
| | 1 vs. 3 | *** | 0.0004 |
| | 1 vs. 4 | **** | <0.0001 |
| WT Female PTX | 1 vs. 2 | ** | 0.0072 |
| | 1 vs. 3 | ** | 0.0049 |
| | 1 vs. 4 | **** | <0.0001 |
| cHET Male PTX | 1 vs. 2 | *** | 0.0004 |
| | 1 vs. 3 | **** | <0.0001 |
| | 1 vs. 4 | **** | <0.0001 |
| WT Male PTX | 1 vs. 2 | * | 0.0125 |
| | 1 vs. 3 | *** | 0.0009 |
| | 1 vs. 4 | **** | <0.0001 |

**Table S1.** Repeated measures mixed-effects model (REML) with Dunnett's multiple comparison test for PTX-treated female and male WT and cHET for TPP test shown in **Fig. 8**.

| **Strain** | **Common Name** | **Strain No.** |
|---|---|---|
| C57BL6/J | B6 | #000664 |
| B6.129X1-*H2-Ab1*$^{tm1Koni}$/J | MHCII$^{fl/fl}$ | #013181 |
| B6.129-*Trpv1*$^{tm1(cre)Bbm}$/J | TRPV1$^{Cre}$ progenitor line | #017769 |
| B6.Cg-*Gt(ROSA)26Sor*$^{tm14(CAG\text{-}tdTomato)Hze}$/J | td-Tomato reporter line | #007914 |

**Table S2.** Jackson Laboratory mouse strains used in the study.

| **Gene** | **Primer Name** | **Sequence** |
|---|---|---|
| MHCII | common forward | 5'-CTC TAC ACC CCC AAC ACA CC-3' |
| | wildtype reverse | 5'-AGT GAG CGA GCA CAG ACA AG-3' |
| | floxed reverse | 5'-TCG CCT TCT TGA CGA GTT CT-3' |
| CRE | Forward | 5'-TTC CCG CAG AAC CTG AAG ATG-3' |
| | Reverse | 5'-CCC CAG AAA TGC CAG ATT ACG-3' |

**Supplemental Table 3.** Primer sequences for TRPV-1 lineage MHCII conditional knockout mice.

| Antibody | Vendor | Clone | RRID | Dilution (application) |
|---|---|---|---|---|
| UCHL1/PGP9.5 | Proteintech | polyclonal | AB_2210497 | 1:12500 (IHC); 1:15000 (ICC); 1:114 (FC) |
| CD3 | Bio-legend | 145-2C11 | AB_312667 | 1:100 (IHC) |
| CD4 | Bio Xcell | clone GK1.5 | AB_1107636 | 2 µg/ml (IHC) |
| MHC Class II, I-A/I-E | BioLegend | M5/114.15.2 | AB_313329 (APC) AB_493525 (647) AB_493523 (488) | 1:2500 (IHC, ICC); 1:100 (FC); 3 µg (western) |
| Rat IgG2b kappa | eBioscience (APC) Biolegend (AF647) | polyclonal | AB_470176 (APC) AB_389343 (AF647) | 1:2500 (IHC, ICC); 1:100 (FC) |
| GLAST | Miltenyi | ACSA-1 | AB_2811532 | 1:50 (IHC) |
| FABP7 | Invitrogen | polyclonal | AB_2542449 | 1:5000 (IHC) |
| beta-tubulin | Abcam | polyclonal | AB_869991 | 1:8000 (western) |
| CD11b 488 | Biolegend | M1/70 | AB_389305 | 1:100 (IHC) |
| CD11c 488 | Biolegend | N418 | AB_389306 | 1:100 (IHC) |
| rabbit DyLight405 | JIR | polyclonal | AB_2340616 | 1:200 (IHC) |
| Armenian hamster 488 | JIR | polyclonal | AB_2338996 | 1:400 (IHC) |
| rat Cy-3 | JIR | polyclonal | AB_2340667 | 1:200 (IHC) |
| rabbit AlexaFluor488 | JIR | polyclonal | AB_2313584 | 1:200 (IHC, ICC) |
| Rabbit 647 | JIR | polyclonal | AB_2492288 | 1:200 (IHC); 1:2000 (western) |

**Supplemental Table 4.** Antibody details for each application. JIR= Jackson ImmunoResearch.

**Supplemental Figures**

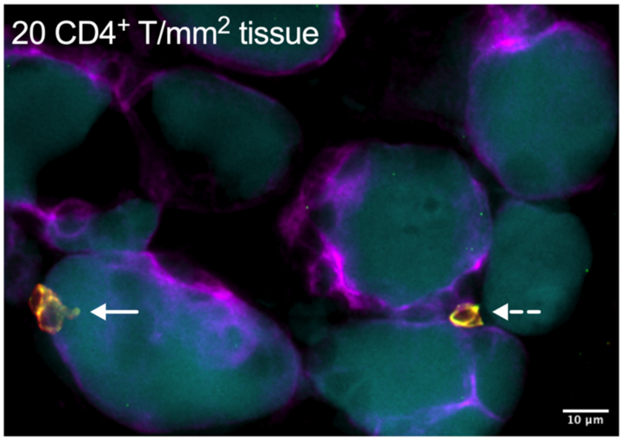


**Fig. S1. CD4$^+$ T cells are found in close proximity to neurons in naïve female DRG tissue.** IHC staining of neurons (PGP9.5$^+$, cyan), SGCs (GLAST$^+$, magenta), and CD4$^+$ T cells (CD3$^+$/CD4$^+$, green/red) in L4 DRG tissue from a naïve female mouse. Representative widefield fluorescence image of CD4$^+$ T cells in close proximity to DRG neurons in naïve female DRG tissue (Solid arrow = absence of GLAST; dashed arrow = occluded by GLAST).

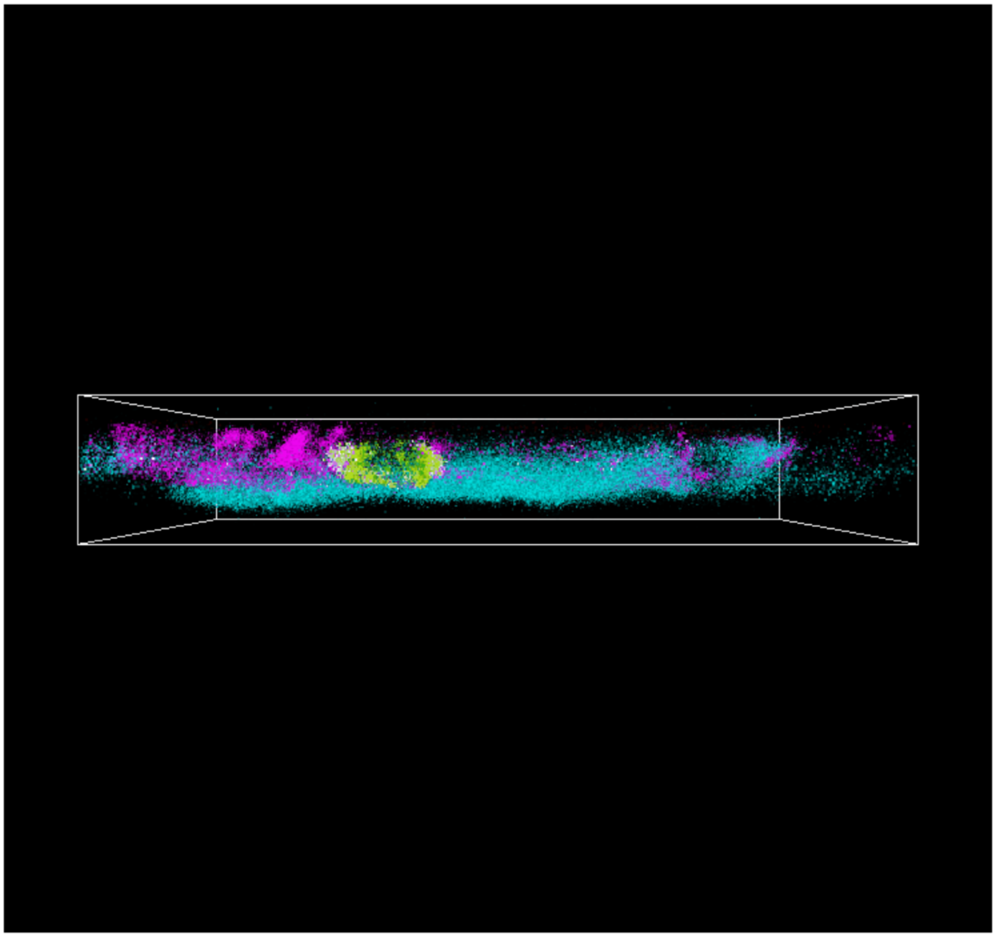

**Fig. S2. CD4$^+$ T cell breaches the SGC barrier in female mouse DRG tissue.** 3D projection of confocal Z-slice images of a CD4$^+$ T cell in direct contact with a DRG neuron from **Fig. 1B, C.** 3D volume rotated around the X-axis at a rate of 10 frames/second (see **FigS1.GIF**).

**A. Day 14 PTX**

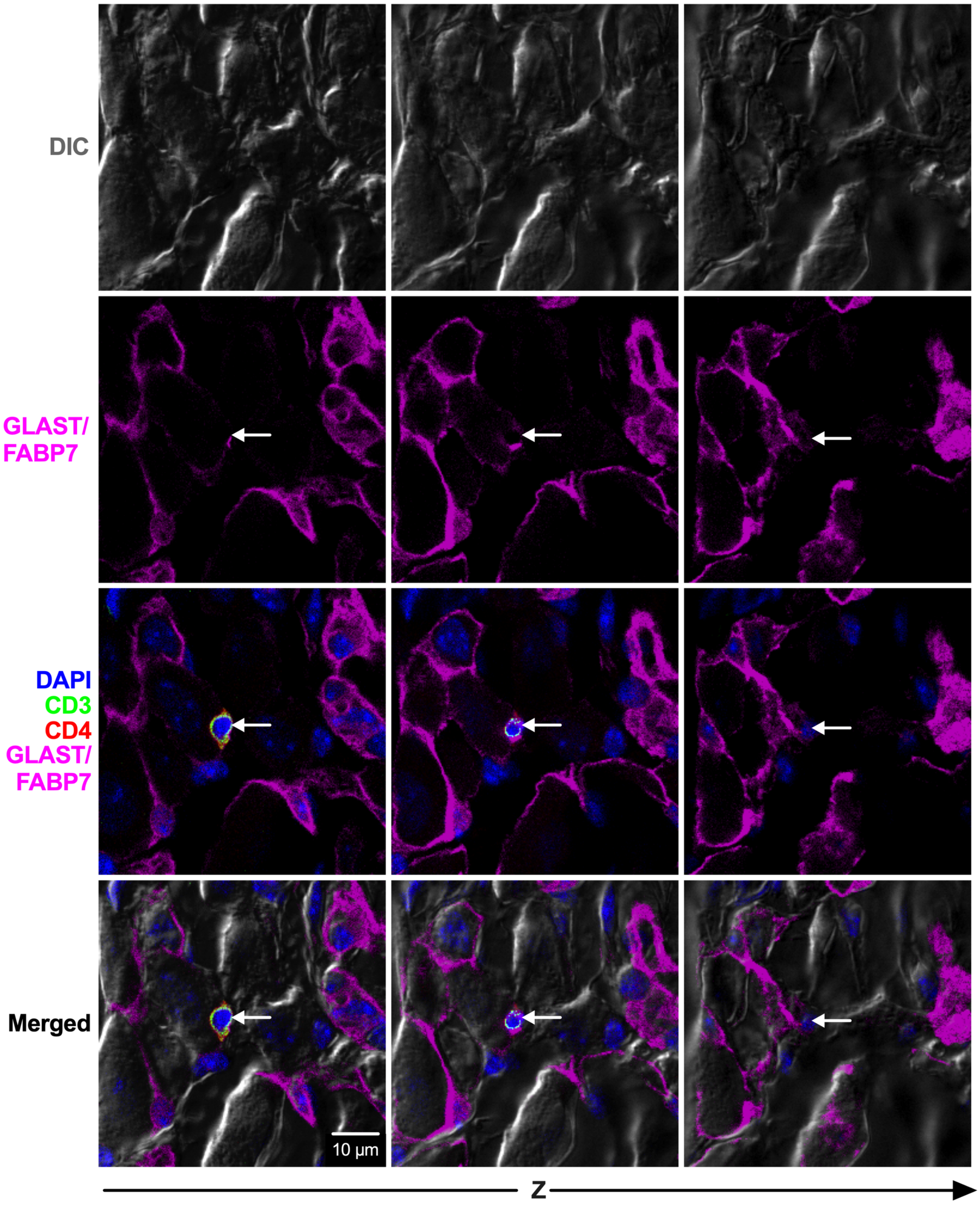


**B. Day 14 PTX**

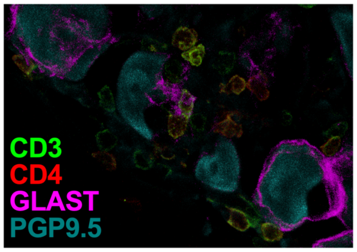

**Fig. S3. CD4$^+$ T cells are in close proximity to neurons in female mouse DRG tissue. (A)** Differential inference contrast (DIC), fluorescence, and merged confocal Z-slice images of a CD4$^+$ T cell in close proximity to a DRG neuron. **(B)** Hotspots of CD4$^+$ T cells in the DRG of a day 14 PTX-treated female mouse.

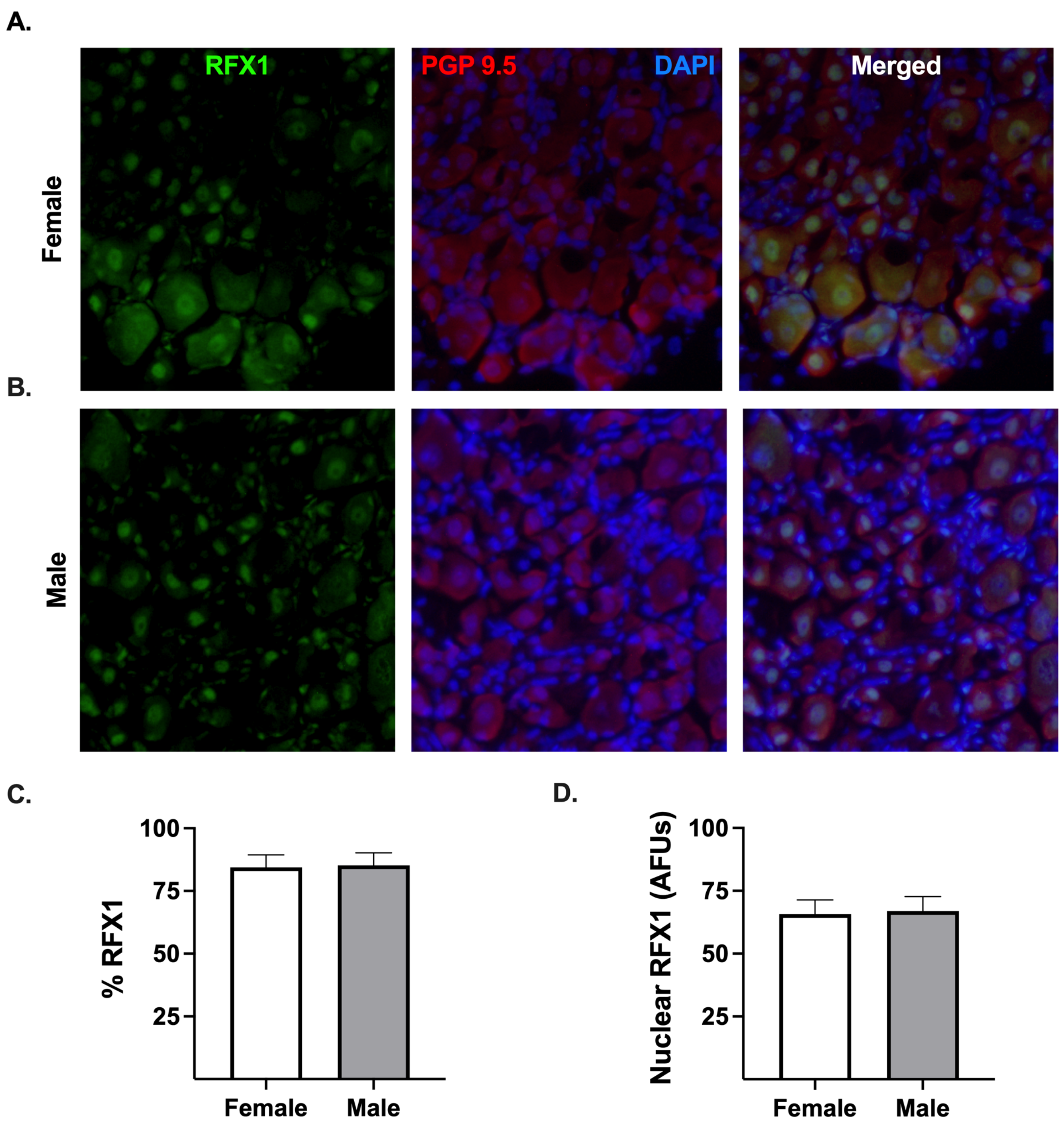


**Fig. S4. DRG neurons from naive female and male mice express nuclear RFX1 protein.** (**A, B**) Immunohistochemistry of RFX1 (green) staining in the nucleus (DAPI, blue) and cytoplasm of DRG neurons ($PGP9.5^{+}$, red) from naïve female (**A**) and male (**B**) mice. (**C**) Percent of $RFX1^{+}$ DRG neurons and (**D**) nuclear neuron RFX1 intensity (AFUs) for naïve female (white bar) and male (gray bar) mice. Statistical analysis by unpaired t-test, n = 8/treatment.

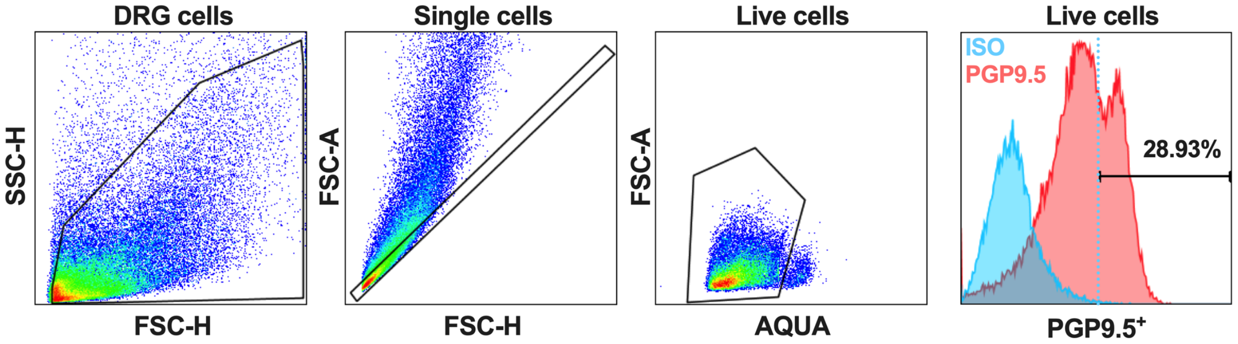


**Fig. S5. PGP9.5$^{+}$ neurons comprise a fraction of total DRG cells.** A nested gating strategy was used to identify PGP9.5$^{+}$ neurons from total DRG cells. Representative histogram overlay of PGP9.5 (red histogram) and Rat IgG2b kappa Isotype Control (ISO; blue histogram with dashed line representing the negative/positive cutoff).

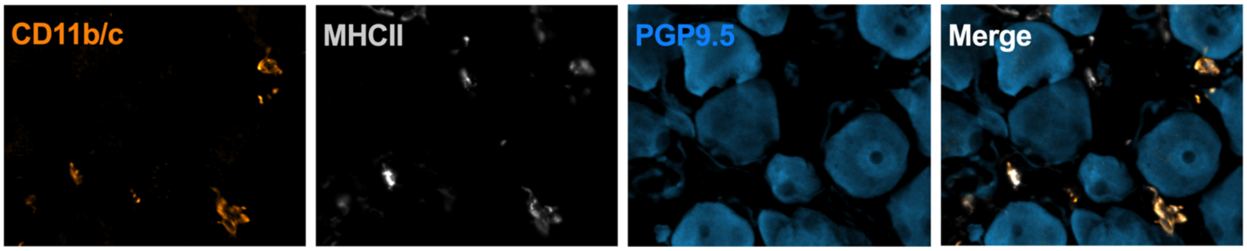


**Fig. S6. CD11b/c$^{+}$ immune cells in DRG tissue express MHCII.** IHC staining of CD11b and/or CD11c (orange), MHCII (gray), and PGP9.5 (blue) in naïve male L4 DRG tissue. Representative single channel and merged widefield fluorescence images of immune cells in DRG tissue co-stained with CD11b/c and MHCII.

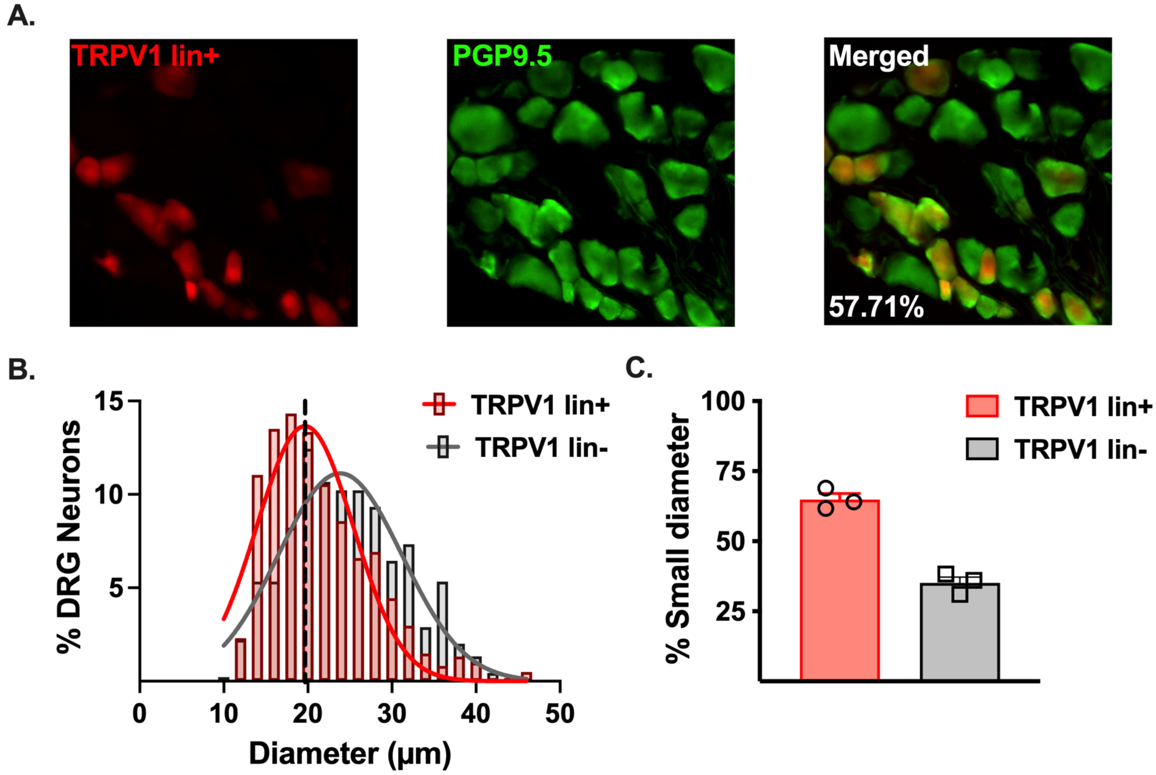


**Fig. S7. High throughput automated image analysis module identifies small and large diameter neurons in female and male DRG tissue.** (**A**) Representative image of td-Tomato-reporter in TRPV1-lineage (red) neurons in naïve female DRG. Percent of TRPV1-lineage neurons from total PGP9.5$^{+}$ (green) neurons shown in merged. (**B**) Gaussian distribution of the diameter (μm) of TRPV1-lineage (red) and non-TRPV1-lineage (gray) neurons. Black dashed line: mode of diameter of TRPV1-lineage neurons (19.66 μm). **(C)** Percent of small diameter TRPV1-lineage positive (red bar) and negative (gray bar) neurons, n=3.

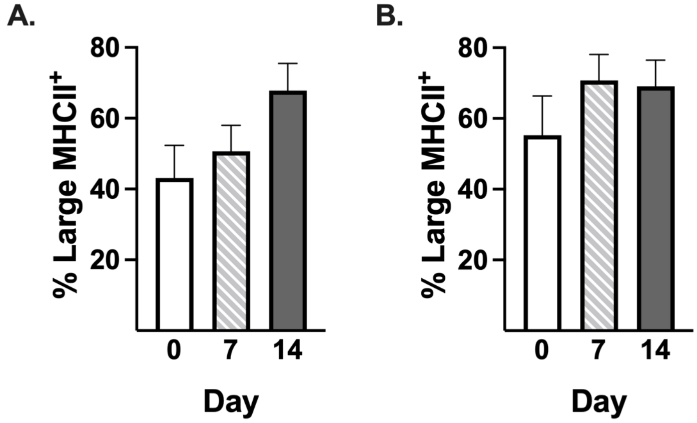


**Fig. S8. PTX does not change the percent of large diameter neurons that express MHCII in male and female DRG.** Percent of large diameter (≥30 μm) neurons positive for MHCII in female (**A**) and male (**B**) DRG before (day 0) and after PTX treatment (day 7 and 14), n=8/treatment.
